# Multi-omic analysis of the cardiac cellulome defines a vascular contribution to cardiac diastolic dysfunction in obese female mice

**DOI:** 10.1101/2022.03.24.485542

**Authors:** Malathi S. I. Dona, Ian Hsu, Alex I. Meuth, Scott M. Brown, Chastidy Bailey, Christian G. Aragonez, Bysani Chandrasekar, Luis A. Martinez-Lemus, Vincent G. DeMarco, Laurel A. Grisanti, Iris Z. Jaffe, Alexander R. Pinto, Shawn B. Bender

## Abstract

Coronary microvascular dysfunction (CMD) is associated with cardiac dysfunction and predictive of cardiac mortality in obesity, especially in females. Emerging evidence suggests development of heart failure with preserved ejection fraction in females with CMD and that mineralocorticoid receptor (MR) antagonism may be more efficacious in obese female, versus male, HFpEF patients. Accordingly, we examined the hypothesis that smooth muscle cell (SMC)-specific MR deletion prevents obesity-associated coronary and cardiac diastolic dysfunction in females. Obesity was induced in female mice via western diet (WD) feeding alongside littermates fed standard diet. Initial studies revealed that global MR blockade with spironolactone prevented impaired coronary vasodilation and diastolic dysfunction in obese females. Importantly, specific deletion of SMC-MR similarly prevented obesity-associated coronary and cardiac dysfunction. Cardiac gene expression profiling suggested reduced cardiac inflammation in WD-fed mice with SMC-MR deletion independent of blood pressure, aortic stiffening, and cardiac hypertrophy. Further mechanistic studies utilizing single-cell RNA sequencing of non-cardiomyocyte cell populations revealed novel impacts of SMC-MR deletion on the cardiac cellulome in obese mice. Specifically, WD feeding induced inflammatory gene signatures in multiple non-myocyte populations (B/T cells, macrophages, and endothelium), independent of cardiac fibrosis, that was prevented by SMC-MR deletion. Further, SMC-MR deletion induced a basal reduction in cardiac mast cells and prevented WD-induced cardiac pro-inflammatory chemokine expression and leukocyte recruitment. These data reveal a central role for SMC-MR signaling in obesity-associated coronary and cardiac dysfunction thus supporting the emerging paradigm of a vascular origin of cardiac dysfunction in obesity.

## Introduction

Coronary microvascular dysfunction (CMD) is independently predictive of cardiac morbidity and mortality in obesity and diabetes^1^. Importantly, females are more likely than males to develop CMD rather than obstructive coronary artery disease^2^. While premenopausal women are protected from heart disease relative to men, that protection is lost in females with obesity and metabolic dysfunction^3^. A recent report revealed close association of CMD and cardiac diastolic dysfunction in females, but not males, with obesity and diabetes^4^. This association has significant implications for ultimate outcomes since patients with both CMD and diastolic dysfunction have a >5-fold increased risk of heart failure with preserved ejection fraction (HFpEF) hospitalization versus patients with isolated CMD or diastolic dysfunction^5^. Accordingly, it has been suggested that co-morbidity-associated CMD may be a primary mechanism of cardiac diastolic dysfunction and HFpEF^6, 7^, conditions for which treatments are limited. While cardiovascular disease mortality is declining in males, it is rising in middle-aged premenopausal females^8, 9^, thus delineation of mechanisms of CMD in females with obesity and metabolic disease is a critical unmet medical need^10^.

Recent evidence, from us and others, revealed that mineralocorticoid receptor (MR) blockade attenuates obesity and diabetes-associated CMD in preclinical models^11, 12^ and patients^13, 14^ independent of blood pressure. Further, pharmacologic MR antagonism with spironolactone (Spiro) prevented obesity-associated diastolic dysfunction in female mice^15^ consistent with a recent *post hoc* analysis of the Treatment of Preserved Cardiac Function Heart Failure with an Aldosterone Antagonist (TOPCAT) trial suggesting greater benefit of MR blockade in obese female HFpEF patients compared to males^16^. Mechanistically, SMC dysfunction (*i*.*e.*, hypercontractility) has been found to precede endothelial dysfunction in obesity^17^ suggesting a potential role for SMC-MR signaling to contribute to impaired coronary and cardiac function in obese females. Indeed, deletion of SMC-MR abrogates aging-associated arteriolar hypercontractility and hypertension^18, 19^. The role of SMC-MR in obesity-associated coronary and cardiac dysfunction has not been explored, nor has the role of SMC-MR been tested in females. To that end, we hypothesized that SMC-MR deletion would prevent both CMD and cardiac diastolic dysfunction in obese female mice thereby expanding our mechanistic understanding of these common conditions.

To test this hypothesis, we induced obesity in female SMC-MR knockout (SMC-MR-KO) and MR-Intact littermate mice by feeding a high-fat/high-carbohydrate western diet (WD) for 16 weeks. Additional studies were conducted in obese mice treated with and without Spiro. We quantified cardiac diastolic function and characterized the coronary and cardiac phenotype compared to SMC-MR-KO and MR-Intact mice fed control diet. We further determined changes in the cardiac gene expression profile at 16 weeks of feeding and utilized single cell RNA sequencing (scRNA-seq) to determine gene expression changes in the non-cardiomyocyte fraction of the mouse heart in response to WD in the presence or absence of SMC-MR. We show that SMC-MR deletion protects females from obesity-induced cardiac and coronary dysfunction, scRNA-seq analysis revealed novel cell-specific gene expression changes induced by obesity that are modified by SMC-MR. These findings support the emerging concept of a vascular origin of cardiac diastolic dysfunction in obesity.

## Methods

Detailed methods available in the Supplemental Material.

## Results

### Systemic MR blockade and SMC-specific MR deletion do not impact traditional cardiac risk factors in obese female mice

WD feeding-induced phenotypic changes in female mice were unchanged by MR blockade with Spiro (Table S1), as we previously reported^15, 20^, nor by SMC-specific MR deletion (Figure 1A, Table S2). Specifically, WD-induced increases of blood glucose, plasma insulin, plasma aldosterone, and plasma cholesterol as well as WD-induced proteinuria and increased urinary blood urea nitrogen (BUN) levels were unchanged by SMC-MR deletion (Table S2). Similar to a recent report^21^, SMC-MR-KO mice fed WD exhibited a modest (∼10%) reduction in average body weight and reduced periovarian adipose weight, compared to MR-Intact controls fed WD (Table S2), with no change in glucose intolerance (Figure S1).

**Figure 1.**
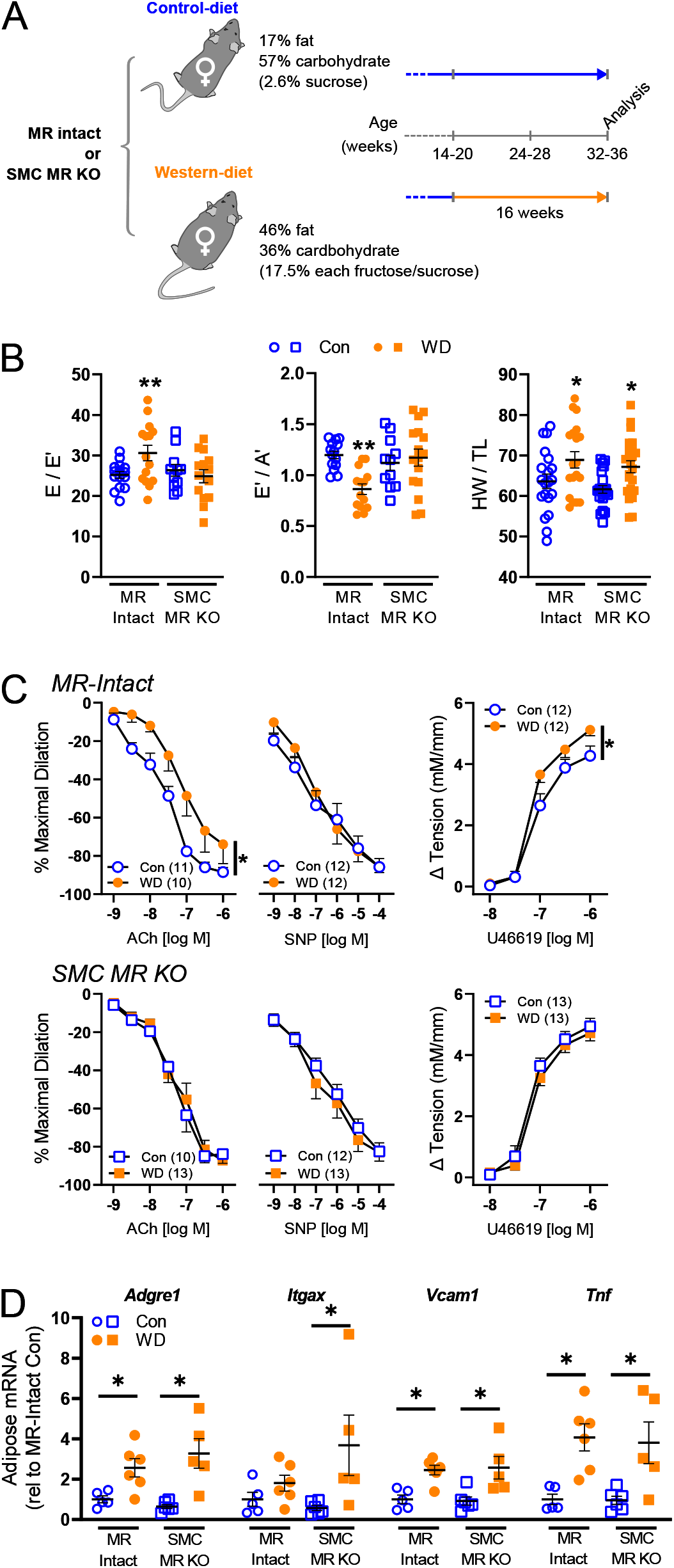
Smooth muscle cell mineralocorticoid receptor knockout (SMC-MR-KO) prevents cardiac diastolic and coronary vascular dysfunction in western diet (WD)-fed female mice independent of adipose inflammation. (A) Mouse cohorts and experimental conditions of the study. All mice were analyzed 16 weeks after Western diet (WD) or control diet (Con) feeding at 32-36 weeks of age. (B) Indices of cardiac diastolic function, specifically estimated left ventricular filling pressure (E/E’) and early-to-late diastolic septal annulus motion ratio (E’/A’), and cardiac weights (heart weight-to-tibia length ratio; HW/TL) in control (Con) and WD fed mice. (C) Vasodilator responses of isolated coronary arteries to endothelium-dependent (acetylcholine, ACh), independent (sodium nitroprusside, SNP) agonists as well as vasoconstrictor responses to the thromboxane A2 analog U46619. (D) Expression of inflammatory genes in reproductive adipose tissue. Values are mean±SE with individual data points shown (A, C). *p<0.05 versus Con or comparison indicated; **p<0.05 versus all other groups.

### Systemic MR blockade and SMC-specific MR deletion prevent WD-induced cardiac diastolic and coronary vascular dysfunction independent of blood pressure and adipose inflammation

WD feeding impaired cardiac diastolic function, indicated by increased LV filling pressure (E/E’), reduced septal wall motion in diastole (E’/A’), left atrial distension (LA/Ao), and induced cardiac hypertrophy in female mice (Figure 1B and S2, Table S3 and S4). Results herein extend prior work by demonstrating concomitant impairment of coronary endothelium-dependent vasodilation with WD feeding in female mice. Furthermore, MR blockade with Spiro attenuated both the coronary endothelial and cardiac dysfunction, but not the cardiac hypertrophy, induced by WD feeding (Figure S2, Table S3). Additional mechanistic studies revealed similar protective effects of SMC-specific MR deletion in female mice (Figure 1). Indeed, unlike WD-fed MR-Intact mice, SMC-MR-KO mice fed a WD did not exhibit impaired diastolic function despite similar WD-induced cardiac hypertrophy (Figure 1B, Table S4). Further, impaired endothelium-dependent vasodilation and enhanced vasoconstriction to the thromboxane analog U46619 induced by WD feeding in MR-Intact mice was absent in WD-fed SMC-MR-KO mice (Figure 1C). There were no differences in diameters of coronary vessels studied (Table S5). Importantly, blood pressure and aortic pulse wave velocity (*i*.*e.*, aortic stiffness) were not changed by either WD feeding or SMC-MR deletion (Table S2). Lastly, WD-induced visceral adipose inflammation indicated by increased gene expression of *Adgre1, Itgax, Vcam1*, and *Tnf* (primers in Table S6), was unchanged by SMC-MR-KO mice (Figure 1D).

### SMC-specific MR deletion shifts WD-induced changes in the cardiac transcriptome

Since SMC-MR deletion did not change traditional risk factors, we explored local cardiac-specific changes associated with SMC-MR signaling in the setting of WD feeding. Specifically, we performed bulk RNA sequencing of LV tissues and examined the unique cardiac transcriptomic signatures induced by WD feeding (Figure 2A; Figure S3) and how the transcriptome differed with SMC-MR deletion in the setting of WD feeding (*i*.*e.*, compared to WD-fed MR-Intact; Figure 2B). Differential gene expression analysis did not reveal any genes/pathways altered as a result of SMC-MR-KO (compared to MR-Intact) in control chow-fed mice (Table S7), however, key pathways that were identified using gene ontology (GO) enrichment analysis across other group comparisons included: water homeostasis (down-regulated in WD-fed versus control-fed MR-Intact mice), circadian regulation (up-regulated in WD-fed SMC-MR-KO WD versus WD-fed MR-Intact mice), and ketone metabolism (down-regulated in WD-fed SMC-MR-KO versus WD-fed MR-Intact mice) (Table S7). Changes in water homeostasis and circadian regulation is consistent with established associations of MR in regulating these biological processes. Moreover, Ingenuity Pathway Analysis (IPA) analysis of differentially expressed genes in WD-fed versus control-fed MR-Intact mice, revealed enrichment of ‘hypertrophy’ (consistent with increased HW/TL with WD feeding) and ‘quantity of reactive oxygen species (ROS)’ biological processes in the top gene network (Figure S4). Accordingly, increased ROS in WD-fed MR-Intact mice was confirmed by restoration of coronary endothelium-dependent vasodilation by the superoxide dismutase mimetic Tempol (Figure S4). Further, in WD-fed SMC-MR-KO compared to WD-fed MR-Intact mice, the top gene network was enriched for biological processes including ‘inflammation of organ’ and ‘leukocyte migration’ (Figure 2C). Interestingly, WD-induced cardiac transcriptomic changes were unique across genotypes with only 3 shared genes (Figure 2D). Directional gene changes in these IPA pathways in WD-fed SMC-MR-KO mice were generally consistent with reduced inflammation and leukocyte migration. Accordingly, assessment of aortic adhesion molecule expression demonstrated upregulation of *Vcam1* and *Icam1* in WD-fed MR-Intact mice indicative of vascular inflammation that is prevented in WD-fed SMC-MR-KO mice (Figure 2E).

**Figure 2.**
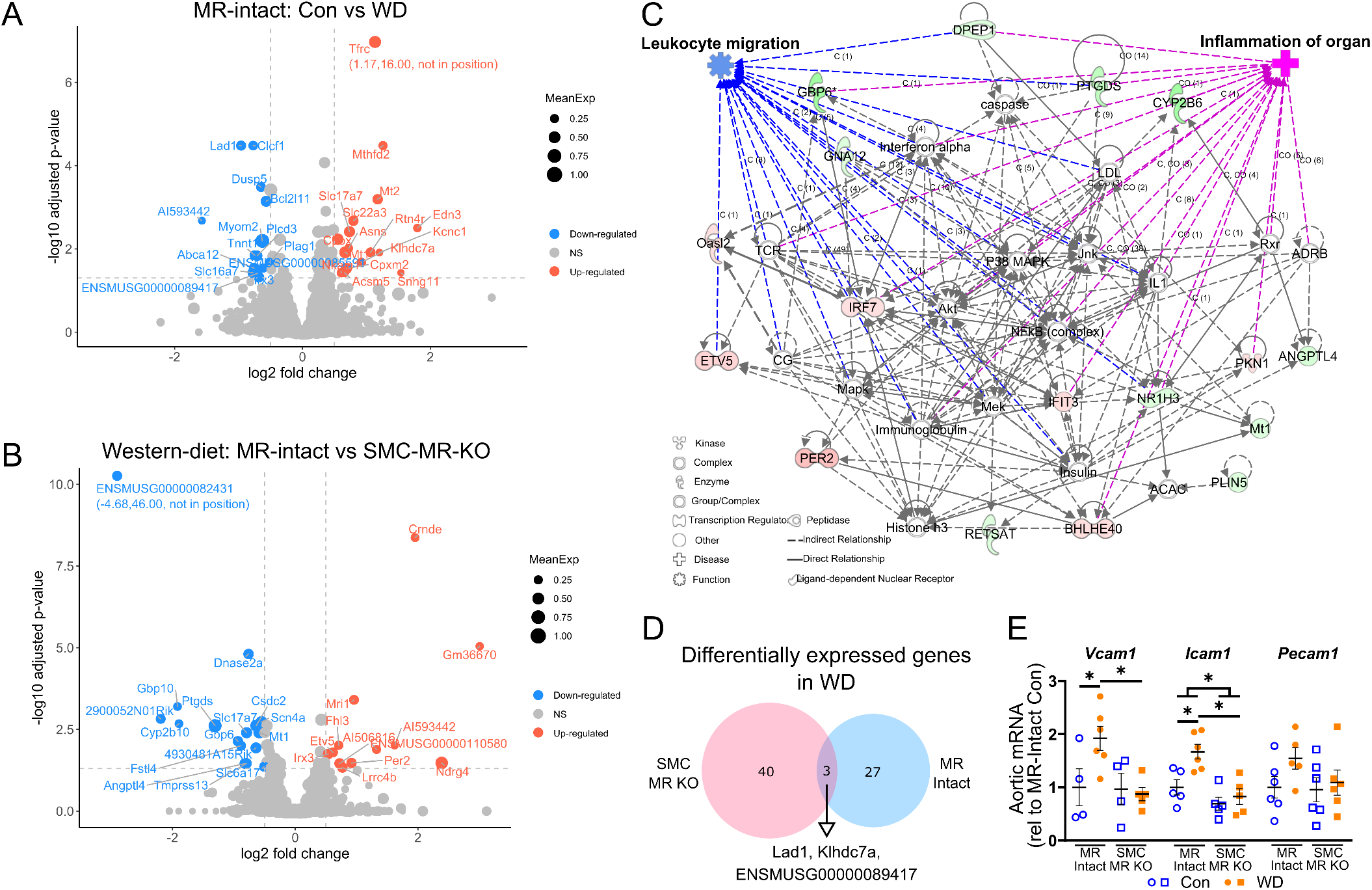
Smooth muscle cell mineralocorticoid receptor knockout (SMC-MR-KO) alters western diet (WD)-induced changes of the cardiac transcriptome. (A) Analysis of cardiac transcripts (13,598 transcripts) revealed differential expression of 30 transcripts induced by WD feeding in MR-Intact mice. Blue and red circles indicated genes down- or up-regulated after WD (14 downregulated, blue dots; 16 upregulated, red dots. (B) Differential gene expression in WD-fed SMC-MR-KO mice versus WD-fed MR-Intact mice (16 downregulated, blue dots; 12 upregulated, red dots). Colored circles in volcano plots (panels A and B) indicate genes with log2 fold change >0.5 and corrected p<0.05. (C) Top differentially regulated IPA network (IPA score=42; green nodes, downregulated; red nodes, upregulated) in WD-fed SMC-MR-KO versus WD-fed MR-Intact mice with enriched relevant biological processes. (D) WD feeding induced unique transcriptomic signatures in WD-fed MR-Intact and SMC-MR-KO mice (3 overlapping differentially expressed [log2 fold change >0.5, corrected p<0.05] transcripts). (E) Gene expression of adhesion molecules in whole aortic tissue from each group. Values are mean±SE with individual data points shown; *p<0.05.

### SMC-MR deletion alters non-myocyte gene signatures in mice fed control diet

Since bulk RNA sequencing analysis is largely dominated by cardiomyocyte mRNA, we next performed high-resolution scRNA-seq analysis of the non-myocyte cell populations to elucidate cell-specific changes elicited by WD feeding and potential mechanisms underlying protection via SMC-MR deletion in obesity. Using established protocols (Figure 3A)^22^, our analysis demonstrated a wide diversity of cardiac non-myocytes including multiple clusters of fibroblasts, vascular smooth muscle and endothelial cells, and diverse immune cell types identified by expression of canonical and non-canonical marker genes (Figure 3B-D, Table S8). The relative proportions of the various cell types did not differ in response to diet or to SMC-MR deletion (Figure S5). We confirmed the SMC-specificity of the model with a reduction of *Nr3c2* (the gene for the MR) across all coronary SMC clusters from SMC-MR-KO compared to MR-Intact mice (Figure 3E-F; Figure S6) with no reduction in MR expression in other non-myocyte cell populations in the heart (Figure S7). Our first analysis compared gene expression between MR-Intact and SMC-MR-KO mice fed control diet. In SMCs, the most differentially expressed gene in response to SMC-MR deletion was a marked up-regulation of the estrogen receptor (*Esr1*) independent of diet feeding (Figure 3G). We also found significant gene expression differences between genotypes in a cell-specific manner (Figure 3H, Table S9). Analysis of differentially regulated genes indicates a key feature of SMC-MR-KO is increased expression of major histocompatibility complex (MHC) class 1 genes (*H2-Q7, -Q4, -Q6, H2-Eb1*, and others) in multiple cell types of SMC-MR-KO animals, particularly ECs, macrophages, and fibroblasts corresponding to GO terms associated with immune modulation and T cell-mediated cell targeting (Figure S8, Table S10). Also noted were increased levels of transcripts corresponding to “response to glucocorticoid” and “response to corticosteroid” in fibroblast and EC subsets, and “angiogenesis” in fibroblasts and SMCs from SMC-MR-KO mice (Table S10). Gene programs corresponding to “angiogenesis” were also frequently down-regulated in multiple cell types, in SMC-MR-KO animals, in addition to those involved in “extracellular matrix organization” (in fibroblast subsets) (Figure S8, Table S10). However, no differences in inflammation, coronary function, or cardiac function were detected relative to control-fed MR-Intact mice (Figures 1 and 2), hence the relevance of these differences in gene expression and programs identified in unstressed mice is unclear.

**Figure 3.**
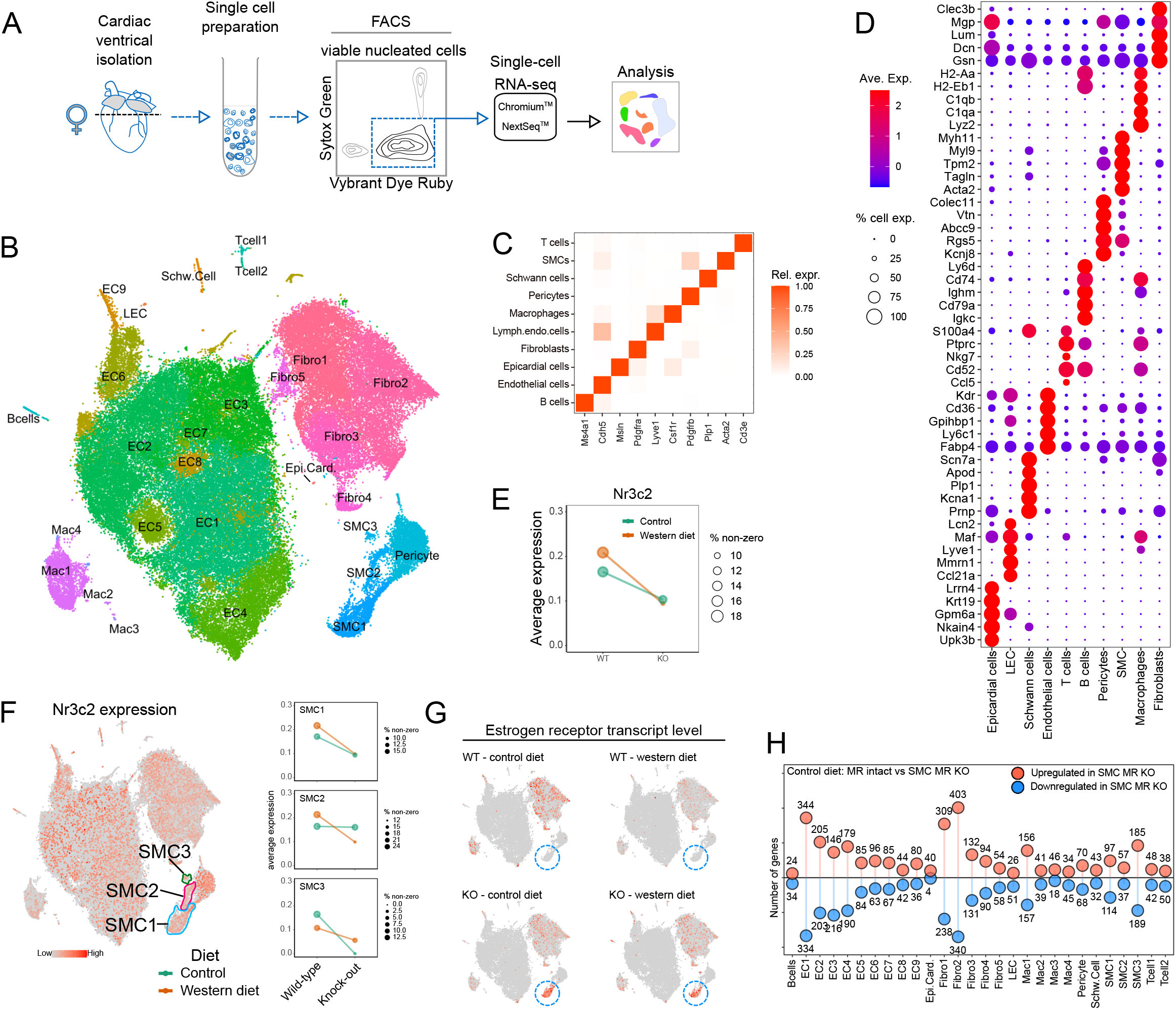
Isolation and analysis of cardiac non-myocyte populations by single-cell RNA-sequencing (scRNA-seq). (A) Schematic outlining the experimental procedure for cell isolation and analysis of adult mouse cardiac non-myocytes by scRNA-seq. (B) t-SNE projection of cardiac cell populations identified by scRNA-seq analysis. Each dot represents a cell that is colored based on distinct cell populations. (C) Heatmap showing relative expression of canonical cell type markers for major cell types identified in adult mouse heart. (D) Dot plot for top 5 highly and uniquely expressed genes in each major cardiac cell population identified using an unsupervised analysis. Dot color and size indicate the relative expression and percentage of cells expressing that gene within each cell population, respectively (also see Table S8). (E) Average expression of mineralocorticoid receptor (MR; Nr3c2) in coronary smooth muscle cells (SMC) from MR-Intact and SMC-MR knockout mice fed control (Con) and western diet (WD). Dot color and size indicate the diet group and the percentage of cells expressing Nr3c2 gene within each group, respectively. (F) MR (Nr3c2) gene expression (red dots) in cell populations and in 3 SMC clusters identified using an unsupervised analysis. Dot color and size (right plots) indicate the diet group and the percentage of cells expressing Nr3c2 gene within each group, respectively. (G) Estrogen receptor (Esr1) expression (red dots) in cell populations from each treatment group with SMC1 population indicated by blue circle. (H) Lollipop plot summarizing number of up- and downregulated genes (uncorrected P<0.01) in Con fed SMC-MR-KO mouse heart cells relative to Con-fed MR-Intact cells (also see Table S9).

### scRNA sequencing of non-myocyte cardiac cell populations reveals that WD induces inflammatory pathways in EC and inflammatory cells independent of fibrosis

We next examined the impact of WD on the cardiac cellulome in MR-Intact mice. This examination of disparate cardiac cell types revealed that WD feeding altered the transcriptional profile of all cell populations examined in MR-Intact mice (Figure 4A, Table S11). Genes up-regulated by WD feeding in MR-Intact mice include genes previously implicated in WD-induced pathology such as *Angptl4* (upregulated in ECs and fibroblasts)^23^, *Cyp1a1* (upregulated in ECs)^24^, *Plin2* (upregulated in ECs, fibroblasts and macrophages)^25^, and *Sgk1* (upregulated in ECs, fibroblasts, pericytes, and B cells)^26^ (Figure 4B). Also up-regulated was *Pparγ* (ECs) which positively regulates many of these genes (*Angptl4, Plin2, Sgk1*) and others (*Fabp4, Cd36, Tsc22d1, Hmox1, Aqp7, Ucp2, Klf4*) that are upregulated in ECs after WD^27^ (Table S11). Conversely, genes down-regulated by WD feeding in MR-Intact mice include genes involved in energy metabolism (*Ckb*, downregulated in EC, fibroblasts, Schwann and T cells, and SMCs), matrix regulation (*Plod1*, downregulated in EC and macrophages), and clearance of advanced glycosylation end products (*Dcxr*, downregulated in fibroblasts, Schwann cells, and SMCs) (Figure S9, Table S11).

**Figure 4.**
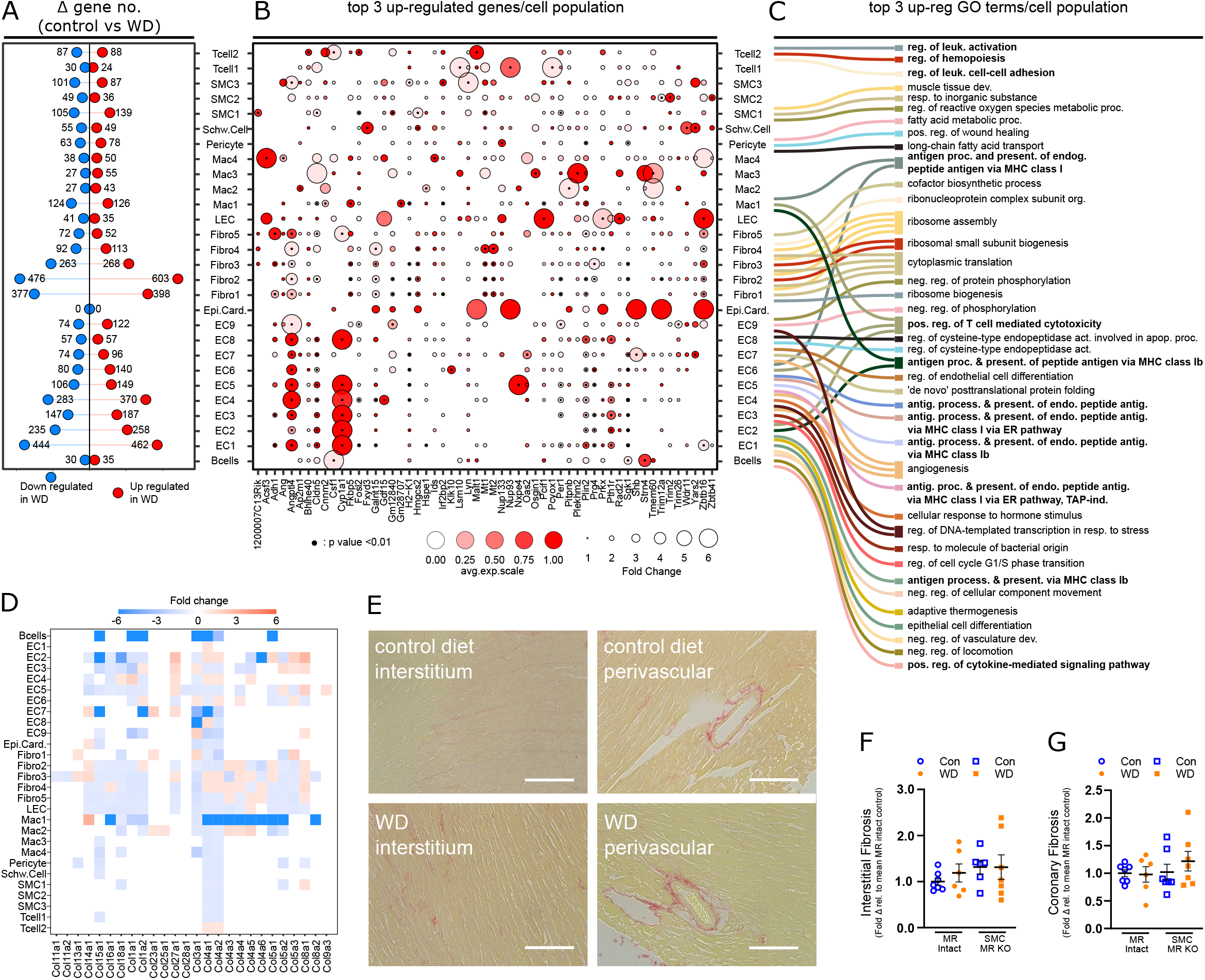
Gene expression changes in cardiac non-myocyte populations in response to western diet (WD) in MR-Intact mice are independent of cardiac fibrosis. (A) Lollipop plot summarizing the number of up- and down-regulated genes (uncorrected p<0.01) identified in WD-fed MR-Intact mice relative to control (Con) diet-fed MR-Intact mice (also see Table S11). (B) Dot plot summarizing the relative expression of top 3 up-regulated genes in response to WD, in each cardiac cell population. Dot color intensity and size are proportional to the relative gene expression in WD cells and the fold change increment in WD cells compared to the Con cells within each cell population, respectively. Black points at the centers of some dots highlight statistically significant differences in gene expression in WD relative to Con cells (uncorrected p<0.01). (C) Sankey plot summarizing the top 3 statistically significant Gene Ontology (GO) terms (corrected p<0.05) enriched by WD up–regulated genes in each cell population. Lines connect GO terms associated with each cell population. Note: not all cell types have 3 statistically significant GO terms (also see Table S12). (D) Heatmap of collagen isoform gene expression in non-myocyte populations from WD versus Con fed MR-Intact mice (see also Figure S12 for all extracellular matrix-related genes). Box color and intensity indicate direction (blue, downregulation; red, upregulation) and magnitude of WD-induced expression changes, respectively. (E) Representative images of cardiac interstitial and perivascular fibrosis assessed by picrosirius red staining. (F-G) Levels of interstitial and coronary fibrosis in mice from all groups, relative to levels in hearts of Con-fed MR-Intact mice. Values are mean±SE with individual data points shown.

To determine genetic programs that are up- and down-regulated following WD feeding we examined GO terms corresponding to biological processes in differentially expressed gene sets (Figure S10). Amongst the top biological processes up-regulated by WD feeding in MR-Intact mice, we found enrichment of terms associated with regulation/function of immune cells (*i*.*e.*, antigen processing and presentation, leukocyte activation) (Figure 4C, Table S12) and that these were also the most commonly up-regulated programs shared across multiple cells populations including ECs and macrophages (Figure 4C). Further, consistent with WD, multiple cell populations also activated gene programs linked to fatty acid and lipid metabolisms and transport (Table S12). Angiogenesis gene programs were both up- and down-regulated by WD feeding (Figure S10, Table S12) in line with no change in capillary density across groups (Figure S11). Lastly, WD feeding did not induce extracellular matrix genes (Figure S12), including collagens (Figure 4D), in non-myocyte populations from MR-Intact mice and GO terms associated with fibrosis (“extracellular matrix organization” and “extracellular structure organization”) were downregulated in major fibroblast clusters (Figure S9, Table S12). Picrosirius red staining confirmed no change in interstitial or perivascular collagen deposition (*i*.*e.*, fibrosis) across all treatment groups (Figure 4E, 4F, and 4G; Figure S12).

### Diabetes- and obesity-associated gene programs are induced by WD feeding independent of MR genotype

Next, we examined the key common gene expression features induced by WD feeding in MR-Intact and SMC-MR-KO mouse hearts. Indeed, a majority of genes altered by WD were equivalently up- or down-regulated across genotypes (Figure 5A-B; Figure S13). Top down-regulated genes included those previously associated with diabetes and obesity, such as *Manf*^28^ and *Creld2*^29^ that facilitate protein folding (Figure 5C), and *Igfbp3* and *Igfbp7*, two structurally similar proteins that regulate the bioavailability of IGFs and insulin. As noted for WD-fed MR-Intact mice, WD also down-regulated genes associated with extracellular matrix in SMC-MR-KO mice (Figure 5C, Table S13 and 14) and top up-regulated genes also included those implicated in diabetes and obesity, such as *Cd36, Txnip, Angptl4, Cyp1a1*, and *Fabp4* (Figure 5D). Consistent with a change in diet, genes corresponding to programs involved in fatty acid and lipid metabolism were up-regulated in both genotypes by WD feeding, however these were primarily restricted to mural cells (Figure 5D, Table S13 and 14). Since SMC-MR-KO mice are protected from WD-induced coronary and cardiac dysfunction, these genotype-independent WD-induced gene changes likely represent pathways not involved in this protection.

**Figure 5.**
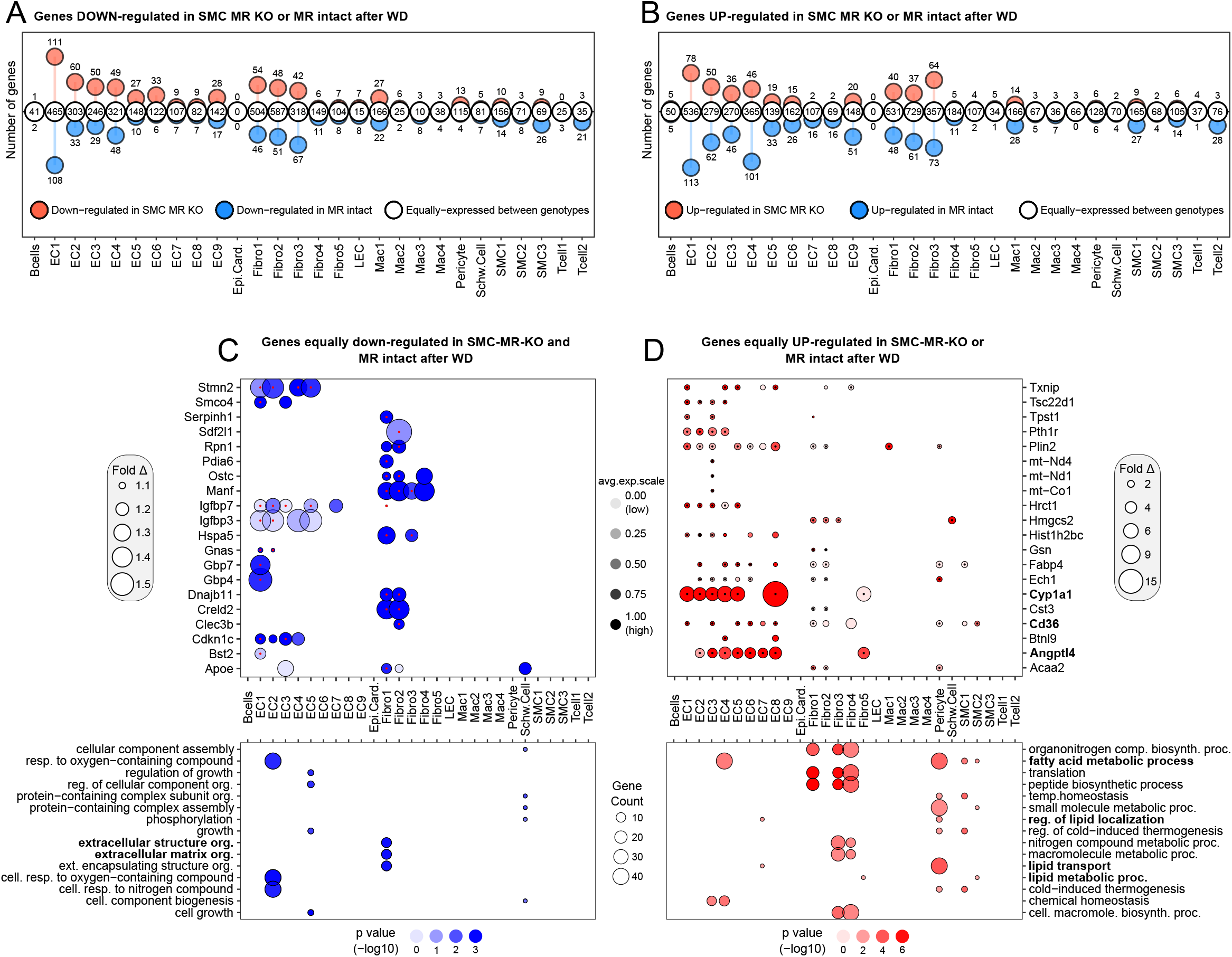
Western diet (WD) feeding induces common and distinct gene activation in MR-Intact and SMC-MR-KO mouse heart cells. (A-B) Lollipop plot summarizing genes similarly (white circles) and differentially down-regulated (A) or up-regulated (B) by WD feeding in MR-Intact (blue circles) and SMC-MR-KO (red circles) mice by cell population. (C-D) Top 20 genes down- or up-regulated (C and D, respectively) following WD in SMC-MR-KO and MR-Intact mice (also see Table S13). Bolded genes indicate those that have been associated with diabetes or obesity. Bottom panels summarize GO terms corresponding to down or up-regulated genes (also see Table S14).

### SMC MR deletion impacts WD-induced gene expression across cell populations, including MR target genes

Next, we examined genes and gene programs that were more robustly induced or repressed by WD feeding in SMC-MR-KO mice compared to MR-Intact mice. Top genes differentially induced or suppressed between genotypes after WD generally followed a cell-specific pattern (Figure 6). Notably, genes whose expression was increased more robustly in MR-Intact mouse hearts included the anti-proliferation genes *Btg3* (ECs and fibroblasts) and *Cdkn1a* (ECs and SMC1), as well as angiogenic genes *Cyr61* (ECs) and *Nrarp* (ECs), and the cholesterol uptake regulator *Ldlr* (fibroblasts and pericytes). Further, genes more robustly expressed in MR-Intact mouse hearts after WD feeding included MR- and GR-sensitive genes such as *Zbtb16, Fam46b*, and *Angptl4* in SMC populations^30^. Closer examination of reported MR-sensitive genes showed up-regulation of a number of these genes in SMCs from MR-Intact, but not SMC-MR-KO, mouse hearts after WD (Figure 6B). Indeed, *Zbtb16*, a key transcriptional repressor, and *Pgf*, a pro-inflammatory MR target gene, were up-regulated by WD in MR-Intact SMCs and down-regulated or not changed, respectively, in SMC-MR-KO SMCs after WD.

**Figure 6.**
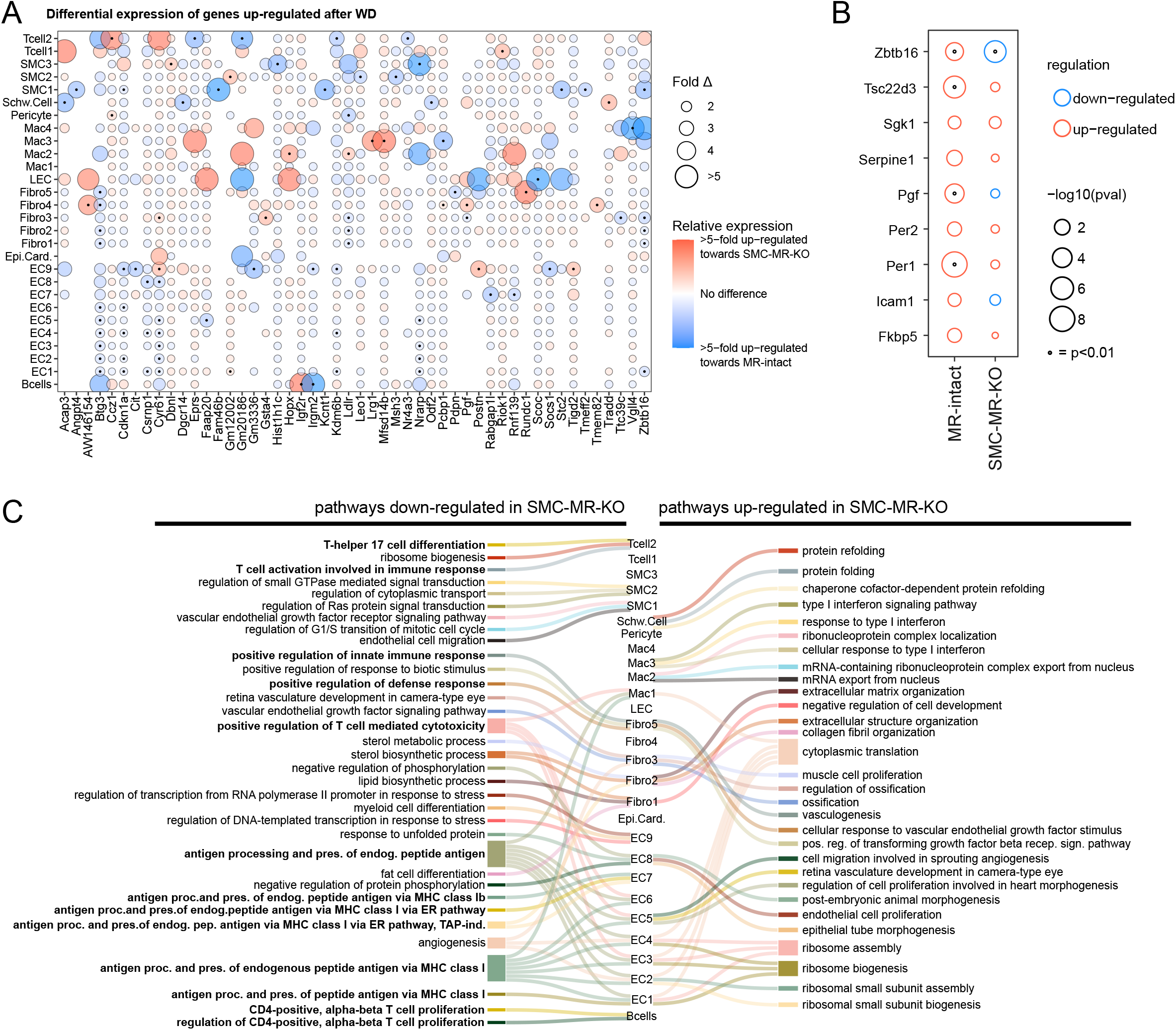
Smooth muscle cell mineralocorticoid receptor knockout (SMC-MR-KO) alters cardiac gene expression response to western diet (WD) feeding. (A) Top-50 genes that are up-regulated after WD and differentially expressed between MR-Intact and SMC-MR-KO mouse heart cells. Blue and red circles indicate genes that are more highly expressed in MR-Intact or SMC-MR-KO mouse heart cells, respectively. Black dot (center of some circles) indicates statistical significance of p< 0.001 (see Table S15). (B) Changes in expression of MR target genes in cardiac SMCs from MR-Intact and SMC-MR-KO mice fed WD versus Con-fed mice from each genotype. Black dot (center of some circles) indicates statistical significance. (C) Sankey plot summarizing the top 3 statistically significant Gene Ontology (GO) terms (corrected p<0.05) enriched in up- (right) and down-regulated (left) genes in WD-fed SMC-MR-KO mice versus WD-fed MR-Intact mice. Lines connect GO terms associated with each cell population. Note: not all cell types have 3 statistically significant GO terms (see Table S16).

### SMC-MR deletion attenuates WD-induced inflammation across the cardiac cellulome

The most significant differences in gene expression programs activated by WD in the two genotypes were those related to immune activity. These were primarily driven by higher expression of genes implicated in inflammation (*Ier2, Iigp1, Egr1, Cyr61, Dusp1, Nr4a1*) and antigen presentation/processing (*Tap1, H2 - D1, -K1, -Q4, -Q6, -Q7, and –T22*) in ECs of WD-fed MR-Intact mouse hearts compared to WD-fed SMC-MR-KO mouse hearts. Intriguingly, in control diet fed mice, many of these genes were more highly expressed in SMC-MR-KO heart ECs, compared to MR-Intact heart ECs (Table S9). Direct comparison of genetic programs among genes down- and up-regulated in SMC-MR-KO mice by WD, versus WD-fed MR-Intact mice, further revealed reduced WD-induced immune response in these mice (Figure 6C; Table S16). Specifically, down-regulated GO terms in WD-fed SMC-MR-KO hearts were enriched for terms associated with antigen processing and presentation (ECs and Mac1) as well as positive regulation of immune responses (ECs, Mac1, Fibro5). Further, down-regulation of GO terms related specifically to T cell immunity were enriched in B cells (*‘*CD4-positive, alpha-beta T cell proliferation’), ECs and Mac1 (‘positive regulation of T cell mediated cytotoxicity’), and T cell2 (‘T cell activation’ and ‘differentiation’) (Figure 6C; Table S14). The enrichment of genes related to leukocyte trafficking and inflammation amongst the most statistically significant programs in multiple cell types (Figure 6C) identifies a key distinction in the phenotypes of MR-Intact and SMC-MR-KO mouse hearts after WD.

Next, using orthogonal approaches, we sought to validate differences in inflammatory responses in the two genotypes identified by scRNA-seq. First, a cytokine array confirmed a pro-inflammatory cytokine signature in hearts from WD-fed MR-Intact mice. Notably, WD feeding upregulated the T cell chemokine lymphotactin and the adhesion molecule L-selectin (Figure 7A). In addition, factors implicated in promotion of local immune/cytokine signaling including MCSF, GITR L, and Axl were upregulated by WD feeding in MR-Intact hearts (Figure 7A). Deletion of SMC-MR altered the cardiac cytokine profile in both control (Figure S14) and WD-fed SMC-MR-KO mice (Figure 7B). In contrast to WD-fed MR-Intact mice, SMC-MR-KO mice fed a WD exhibited cytokine changes indicative of reduced cardiac inflammation compared to control fed SMC-MR-KO mice. Specifically, the pro-inflammatory mediators TWEAK, PDGF-AA, MIP-1b, IL-22, and IL-1a were reduced in WD-fed SMC-MR-KO mice (Figure 7B). Accordingly, the impact of SMC-MR deletion on WD-induced cardiac immunity was further explored by immunohistochemistry and flow cytometry. In MR-Intact mice, WD feeding increased cardiac CD3+ T cells but not CD68+ monocytes/macrophages (Figure 7C and 7D, Figure S15 and S16). Single-cell transcriptomics revealed upregulation of macrophage activation markers *Cd86* and *Cd83* (Mac1; Table S9) in WD-fed MR-Intact mice consistent with a trend for increased cardiac CD11b+CD80+ macrophages by flow cytometry (p=0.07; Figure S16). These changes in cardiac leukocytes were prevented in WD-fed SMC-MR-KO mice. Furthermore, consistent with downregulation of the ‘Th17 cell differentiation’ GO term in the Tcell2 cluster from WD-fed SMC-MR-KO mice compared to WD-fed MR-Intact mice (Figure 6C), SMC-MR deletion was associated with reduced cardiac Th17 cells (Figure S16). Lastly, SMC-MR-KO mice exhibited a reduction in cardiac mast cells, a cell population recently implicated in diabetes-associated cardiac dysfunction^31^, independent of diet feeding (Figure 7E, Figure S16). Together, these data implicate SMC-MR signaling as a critical driver of obesity-associated inflammation across the cardiac cellulome underlying subsequent development of coronary and cardiac dysfunction in obesity.

**Figure 7.**
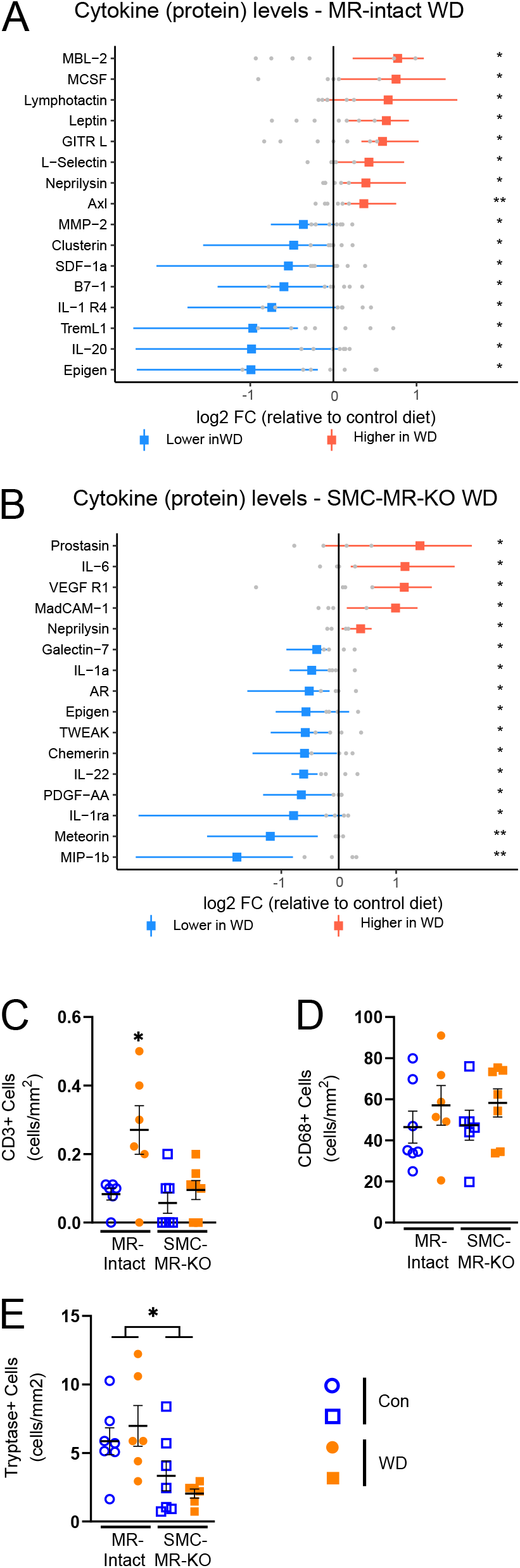
Smooth muscle cell mineralocorticoid receptor knockout prevents obesity-associated cardiac inflammation. Differentially expressed cardiac cytokines in (A) WD-fed MR- Intact mice and (B) WD-fed SMC-MR-KO mice versus Con-fed mice of each genotype. Red, upregulation; blue, downregulation; colored dot is mean with bar indicating spread of data in WD group; gray dots indicate individual data points in Con group. *p<0.05; **p<0.01. Cardiac (C) CD3+, (D) CD68+, and (E) Tryptase+ cells assessed by immunohistochemistry. Values are mean±SE with individual data points shown. *p<0.05 versus all other groups or noted comparison.

## Discussion

Our data demonstrate critical involvement of SMC-MR in obesity-associated coronary and cardiac dysfunction in female mice. Specifically, in obese female mice, SMC-MR deletion prevented the decline in coronary endothelium-dependent vasodilation and increase of vasoconstriction as well as impaired cardiac diastolic function, but not cardiac hypertrophy. Importantly, these benefits of SMC-MR deletion occurred independent of changes in blood pressure, aortic stiffening, and obesity-associated adipose inflammation, metabolic dysregulation, and kidney injury. Further, scRNA-seq revealed a distinct inflammatory gene profile in non-myocyte populations from obese mice that was independent of cardiac fibrosis and associated with cardiac leukocyte infiltration. This obesity-associated cardiac inflammatory phenotype was generally prevented by SMC-MR deletion. Together, these data provide unique insight and support the emerging paradigm of a vascular origin of cardiac dysfunction in obese females^1, 6^, patients at high risk for developing HFpEF^4, 5^.

Involvement of MR signaling in obesity-associated cardiovascular morbidity and mortality is supported by both clinical and preclinical work, from us^11, 12, 15, 32, 33^ and others^13, 14, 34^, utilizing MR antagonists. In obese rodent and swine models, MR blockade with spironolactone or eplerenone mitigated endothelial dysfunction and vascular remodeling as well as diastolic dysfunction, cardiac oxidative stress, fibrosis, and inflammation^11, 12, 15, 32-34^. Mechanistically, several prior studies have revealed an important role for EC MR signaling underlying vascular and cardiac dysfunction in obesity. Specifically, deletion of EC MR prevented obesity-associated endothelial dysfunction in aorta and mesenteric arterioles as well as cardiac diastolic dysfunction^20, 35, 36^. Improved vascular function in obesity following EC MR deletion was associated with modulation of reactive oxygen species production/degradation and NO bioavailability^35, 36^ while prevention of cardiac dysfunction was associated with reduced cardiac pro-oxidant and pro-inflammatory signaling^20^. Improved cardiac function in obese female mice with EC MR deletion may also be due, in part, to attenuated obesity-associated aortic stiffening in these mice^35^. Our data expand this previous work by demonstrating, for the first time, a critical role of SMC-MR signaling underlying coronary and cardiac diastolic dysfunction in obese females. That obesity-associated endothelial dysfunction was prevented by SMC-MR deletion supports recent evidence that SMC dysfunction may precede^17^, and contribute to development of, impaired endothelium-dependent vasodilation in obesity. Taken together, these studies highlight novel, independent roles of MR signaling in vascular cells in the pathogenesis of obesity-associated cardiac dysfunction in females. Importantly, the protection afforded by SMC-MR deletion occurs independent of changes in traditional risk factors (blood pressure, glucose intolerance, hypercholesterolemia, kidney injury) and aortic stiffening suggesting local SMC-MR-dependent mechanisms of dysfunction within the cardiac microenvironment.

Systemic and cardiac inflammation have been implicated as requisite contributors in the etiology of obesity-associated cardiac impairments, including HFpEF^6, 37, 38^. Indeed, cardiac inflammation has been suggested to precede and contribute to other common characteristics of cardiac diastolic dysfunction, such as fibrosis and sarcomeric stiffening, in the setting of co-morbid conditions^6, 39^. Consistent with this paradigm, in our hands, WD-fed MR-Intact female mice do not exhibit cardiac fibrosis despite increased cardiac expression of pro-inflammatory cytokines (*e*.*g.*, neprilysin, M-CSF). These data suggest that WD-induced diastolic impairments in this model likely involve sarcomeric stiffening (*i*.*e.*, titin hypophosphorylation)^40^ and/or altered cardiomyocyte calcium handling^41, 42^.

Unbiased scRNA-seq data confirmed upregulation of inflammatory pathways across cardiac non-myocyte populations by WD feeding independent of a pronounced upregulation of extracellular matrix-related genes. These data further highlight a vascular contribution to cardiac inflammation in obesity via upregulation of gene programs for EC antigen presentation and processing via MHC class I molecules in conjunction with upregulation of genes associated with T cell-mediated cytotoxicity in cardiac ECs and macrophages. These gene changes correspond to cardiac infiltration of CD3+ T cells supporting recent evidence that increased endothelial MHC class I molecule expression enhances T cell transmigration^43^. Further, cytokine analysis reveals obesity-associated upregulation of the adhesion molecule L-selectin, the T cell chemokine lymphotactin, and the T cell costimulatory ligand GITR-L. Interestingly, GITR-L engagement of T cell GITR has been reported to reduce susceptibility of effector T cells to suppression by T regulatory cells^44^. Prior work has established a link between cardiac T cell infiltration and the development of systolic dysfunction in response to cardiac pressure overload^45^. Our data extend these findings and, to our knowledge, are the first to suggest a role for cardiac T cell infiltration in the inflammatory processes contributing to cardiac diastolic dysfunction in obese female mice.

MR-dependent inflammation has been implicated in a variety of disease states, including obesity. Indeed, global MR blockade reduces obesity-associated adipose^11, 12, 46^, cardiac^11, 12, 15, 46^, and vascular^11,32^ inflammation. Our data delineate a novel role for SMC-MR signaling as a primary contributor to coronary and cardiac, but not adipose, inflammation in obesity. Specifically, obesity-associated upregulation of aortic adhesion molecules and endothelial MHC class I molecules as well as cardiac T cell infiltration were prevented in obese SMC-MR knockout mice. This protective effect of SMC-MR deletion corresponded with generally anti-inflammatory shifts in cardiac cytokines in obese, versus lean, SMC-MR knockout mice and prevention of obesity-associated upregulation of the pro-inflammatory SMC-MR target *Pgf*. Similar prevention of cardiac inflammation by SMC-MR deletion, including prevention of cardiac *Pgf* upregulation, was recently reported in male mice subjected to cardiac pressure overload^47^. Further, cardiac leukocyte infiltration in response to pressure overload consisted entirely of CD3-CD11b+ myeloid cells and was prevented by SMC-MR deletion^47^. In conjunction with the present results, these data argue for significant context-specificity of SMC-MR-dependent mechanisms of cardiac inflammation and leukocyte recruitment. Intriguingly, our data reveal a novel reduction of cardiac mast cells in SMC-MR-KO mice independent of diet treatment. Recent evidence in female *db*/*db* mice demonstrated a critical role for activation/degranulation of resident cardiac mast cells in diabetes-associated cardiac leukocyte infiltration and diastolic dysfunction^31^. Thus, reduced cardiac mast cells may be a unifying mechanism of cardioprotection accounting for reduced cardiac dysfunction in SMC-MR-KO mice following obesity, coronary ligation^48^, aging^49, 50^, and pressure overload^47^. Our use of inducible SMC-MR deletion suggests that any impact of SMC-MR signaling on cardiac mast cells is not of developmental origin and potential SMC-MR-dependent mechanisms of cardiac mast cell recruitment/maturation are warranted.

Mechanistically, our data suggest that enhanced SMC estrogen signaling in SMC-MR-KO mice may contribute to prevention of obesity-associated cardiovascular dysfunction. Indeed, SMC-MR deletion resulted in pronounced upregulation of SMC estrogen receptor (ER; *Esr1*) gene expression. This finding is consistent with recent reports of ER upregulation in macrophages/Kupffer cells^51^ and EC^52^ following cell-specific MR deletion. In the latter study, double deletion of EC MR and ER eliminated the prevention of obesity-associated endothelial dysfunction afforded by EC MR deletion alone^52^. While potential SMC ER-dependent mechanisms of cardioprotection remain unclear in the present study, recent evidence supports paracrine SMC ER signaling as a contributor to endothelial healing/regeneration following vascular injury^53, 54^. Since this study was performed only in female mice, whether SMC ER upregulation might occur in male SMC-MR-KO mice is not known. We focused on female mice in this study in light of the high prevalence of coronary microvascular dysfunction in women with co-morbid conditions and its close association with cardiac dysfunction^4, 5, 10^.

Our study also provides a useful resource for examining pathways that may be important for cardiac remodeling, but not WD-induced diastolic dysfunction. These include genes such as such *Angptl4, Cd36, Cyp1a1* and *Pparγ* and others which are upregulated after WD in both MR-Intact and SMC-MR-KO mouse hearts. Examination of these pathways to abrogate cardiovascular remodeling have been active areas of research for many years and master regulators, such as *Pparγ*, may be key for WD-induced changes in EC phenotypes we have determined here by scRNAs-seq. Further, our proteomic analysis also detected neprilysin—an enzyme that degrades natriuretic peptide and angiotensin II—at higher levels after WD in both genotypes. Indeed, targeting neprilysin while inhibiting the angiotensin receptor improves morbidity and mortality associated with heart failure^55^. However, modulating the activity of these elements that are activated in both MR-Intact and SMC-MR-KO mouse hearts may have limited effect on diastolic impairment of heart function in obesity.

In summary, this study reveals a central role for SMC-MR signaling in the development of obesity-associated coronary and cardiac diastolic dysfunction in female mice. This is the first report, to our knowledge, of obesity-associated transcriptomic changes across the cardiac non-myocyte cellulome. This unbiased approach revealed cardiac inflammation, associated with lymphocyte infiltration, and hypertrophy independent of fibrosis in obese female hearts. SMC-MR deletion mitigated obesity-associated cardiac and coronary inflammation and dysfunction, but not hypertrophy, potentially involving reduced cardiac mast cells and enhanced SMC estrogen signaling that warrant further investigation. These results shed new light on vascular mechanisms of obesity-associated cardiac dysfunction in premenopausal women and provide rationale for further study of MR inhibition and pathways downstream of SMC-MR as sex-specific strategies to treat cardiac and coronary dysfunction, critical contributors to development of HFpEF in obese women.

## Acknowledgements

The authors gratefully acknowledge the assistance of Jacob Russell and Maloree Khan. This work was funded by NIH R01 HL136386 (Bender), NIH R01 HL119290 (Jaffe), the University of Missouri Research Core Facilities Grant Program (Bender), and NHMRC Ideas Grant GNT1188503 (Pinto). The work was also supported by resources and the use of facilities at the Harry S. Truman Memorial Veterans Hospital in Columbia, MO.

