## Supplemental Material for "Multi-omic analysis of the cardiac cellulome defines a vascular contribution to cardiac diastolic dysfunction in obese female mice"

*Running title:* Vascular role in cardiac dysfunction in obesity

***Correspondence:**

Shawn B. Bender, Ph.D.
Harry S. Truman Memorial Veterans' Hospital
and University of Missouri
Biomedical Sciences
E102 Vet Med Bldg
Columbia, MO 65211


Alexander R. Pinto, Ph.D.
Baker Heart and Diabetes Institute
75 Commercial Rd
Prahran, Victoria
Australia 3004


**Methods**

*Animals*. All animal protocols were approved by the Institutional Animal Care and Use Committees of the Harry S Truman Memorial Veterans Hospital and the University of Missouri in compliance with the *Guide for the Care and Use of Laboratory Animals* (National Institutes of Health). All animals were housed in a temperature-controlled room (12:12-h light-dark cycle) and provided ad libitum access to water and either a standard control diet (Con; LabDiet 5008) or a western diet (WD; TestDiet 58Y1 modified) consisting of 46% fat and 36% carbohydrate (17.5% each from sucrose and high fructose corn syrup) for 16 wk beginning at 14-26 wk of age. Two experimental paradigms were utilized. First, female C57BL/6J mice (Jackson Labs) were randomly divided into 3 groups: Con, WD, or WD treated with the MR antagonist spironolactone (Spiro, sc, 0.63 $mg \cdot d^{-1}$; Innovative Research of America) for 16 wk. Con and WD-fed mice received placebo pellets for 16 wk. Secondly, mice with inducible MR deletion specifically in smooth muscle cells (SMC-MR knockout [SMC-MR-KO] mice) were generated, as previously described[1, 2], by crossing floxed MR mice with mice containing a Cre-recombinase-$ER^{T2}$ gene driven by the smooth muscle actin promoter (SMA-Cre-$ER^{T2}$) and activated by tamoxifen. These mice are compared with floxed MR/SMA-Cre-$ER^{T2}$ negative (MR-intact) littermates. Both strains were treated with tamoxifen at 6-8 wk of age, resulting in MR deletion in the Cre positive animals as previously confirmed[1, 2]. SMR MR KO and MR-intact mice were fed Con and WD for 16 wk.

*Plasma and urine parameters*. Twenty-four hour urine collection was performed during the final week of diet feeding as was glucose tolerance testing. Following measurement of fasting blood glucose, mice were injected with glucose (ip, 1 g/kg) and blood glucose subsequently measured at 15, 30, 60, and 120 mins post-injection via glucometer (AlphaTRAK 2, Zoetis). On the day of euthanasia, mice were fasted for 5 hours and anesthetized with isoflurane (2-4% in 100% $O_2$); blood was collected via the inferior vena cava, processed to plasma, and frozen at -80°C. Blood glucose was determined by glucometer. Plasma aldosterone was quantified by radioimmunoassay (Tecan MG13051). All other plasma and urine measures were analyzed at Comparative Clinical Pathology Services (Columbia, MO).

*Echocardiography*. During the final week of diet feeding/treatment, transthoracic echocardiography was performed (Vevo2100, FUJIFILMS, Visualsonics, Toronto) with an MS400 high frequency echo probe at the Small Animal Ultrasound Imaging Center at the Harry S Truman Memorial Veterans Hospital. Under anesthesia (0.75-4% isoflurane in 100% $O_2$), mice were placed on a heated platform to maintain body temperature at 37°C and two-dimensional echocardiograms were performed in the apical four chamber view. Initially, a small sample volume was positioned in the left ventricle (LV) just proximal to the mitral leaflets to acquire early (E) and late (A) diastolic blood flow velocities in pulse wave (PW) Doppler mode. Isovolumic relaxation time (IVRT) was also determined from the PW spectra. B- and M-mode images of the LV and septum in short axis view were acquired at the level of the papillary muscles. Left ventricular anterior and septal wall thicknesses at end diastole (LV SWTd and LV PWTd), luminal diameters (LVIDs and LVIDd), and ejection fraction (EF) were determined offline in M-mode. B-mode images in modified long axis (ascending aortic) view were acquired for determination of left atrial and aortic diameters. Next, Tissue Doppler Imaging (TDI) was performed in the apical four chamber view to acquire early (E') and late (A') septal annular velocities. Parameters were assessed using an average of three beats from three different spectra, and calculations were made in accordance with the American Society of Echocardiography guidelines as well as specific guidelines for rodent echocardiography. All data were acquired and analyzed offline by a single blinded observer.

*Aortic vascular measurements and blood pressure*. During the final week of diet feeding/treatment, *in vivo* aortic stiffness was evaluated in isoflurane-anesthetized mice (1.75% in 100% $O_2$) by pulse-wave velocity (PWV) using Doppler ultrasound (Indus Mouse Doppler System, Webster, TX), as previously described[3]. Briefly, using transit time method, PWV was quantified as the difference in arrival times of a Doppler pulse wave at two locations along the aorta at a fixed distance[4]. The distance between the two locations along the aorta is divided by the difference in arrival times and is expressed in m/s. Velocity waveforms were acquired at the aortic arch followed immediately by measurement at the descending aorta 35 mm distal to the aortic arch. Systolic blood pressure was determined by tail-cuff plethysmography (BP-2000, Visitech) during the final week of treatment, as previously described[5].

*Coronary Vasomotor Function*. Following blood collection under anesthesia (2-4% in 100% $O_2$), mice were perfused via the left ventricle at 100 mmHg with ice-cold physiological salt solution (PSS) comprised of (in mM): 119 NaCl, 4.7KCl, 2.5 $CaCl \cdot 2H_2O$, 1.17 $MgSO_4 \cdot 7H_2O$, 1.18 $KH_2PO_4$, 0.027 EDTA, 25 $NaHCO_3$, and 5.5 glucose (pH 7.4). The heart was subsequently removed and segments of the left coronary artery (~1mm) dissected and mounted on 17 µm stainless steel wires in oxygenated PSS (95% $O_2$-5% $CO_2$) in a small vessel myograph for isometric tension recording (Danish Myo Technology, Aarhus, Denmark), as previously described[6, 7]. Vessel length was quantified after mounting with a calibrated ocular micrometer. Following equilibration and normalization using an established procedure[8]. Vessel viability was confirmed by exposure to 80 mM KCl PSS. Following washing, vasoconstrictor responses to the thromboxane $A_2$ analog U46619 (10 nM – 1 µM) were assessed as well as vasodilator responses to acetylcholine (ACh; 1 nM – 0.1 mM) and sodium nitroprusside (SNP; 1 nM – 0.1 mM) following preconstriction with U46619 (100-300 nM). A subset of vessels were pretreated with the superoxide dismutase mimetic Tempol (1 mM for 20 min) prior to assessment of ACh-induced vasodilation. Vasodilator responses are reported as percent maximal dilation from U46619 preconstriction. Vasoconstrictor responses are reported as developed tension normalized to vessel length (mN/mm). Since this vessel does not develop spontaneous myogenic tone, minimum tension (*i.e.*, maximal dilation) was determined following the normalization procedure.

*RT-PCR*. Aortas and periovarian adipose tissue were homogenized in a Tissue-Lyser (Qiagen) and total RNA was extracted using the Qiagen RNeasy Fibrous (aorta) or Lipid (adipose) Tissue kit and quantified using a Nanodrop spectrophotometer (Thermo Scientific). First-strand cDNA was synthesized from total RNA using the Improm-II reverse transcription kit (Promega) and quantitative real-time PCR was performed using the CFX Connect Real-Time PCR Detection System (Biorad) using target specific primers (Table S6). PCR reactions using iTaq Universal SYBR Green SMX (Biorad), thermal conditions, and melt curve analysis were performed as previously described[5]. GAPDH was used as an internal control gene and messenger RNA (mRNA) expression values were calculated based on cycle thresholds (CTs) via the $2^{\Delta\Delta CT}$ method, where $\Delta CT$ = GAPDH CT – gene of interest CT and are presented normalized to control mice, which were set at 1.

*Immunohistochemistry and staining*. Left ventricular tissue was immersion fixed in 10% buffered formalin, dehydrated in ethanol, paraffin embedded, and sectioned in 5 µm slices, as previously described [CITE]. To evaluate fibrosis, sections were stained with picrosirius red (PR) for determination of cardiac interstitial and periarterial collagen. Images were obtained using an EVOS FL Auto Imaging System and quantified using the thresholding function in ImageJ. Periarterial fibrosis was quantified as the ratio of PR-stained periarterial area to luminal circumference. Interstitial fibrosis was quantified as the percent area of myocardial PR staining. Cardiac capillary density was quantified in FITC-conjugated CD31 (1:50, Novus)-stained cardiac sections. In additional sections, following citrate buffer antigen retrieval, hearts were blocked with 10% FBS in PBS and 0.3% $H_2O_2$ to prevent endogenous peroxide activity. Hearts were incubated with antibodies against CD3 (1:100; Abcam #ab5690), CD68 (1:100; Abcam #ab31630), or mast cell tryptase (1:100; Abcam #ab2378). Washed slides were incubated with the appropriate HRP conjugated secondary antibody. Stained hearts were developed using a DAB Substrate Kit (Thermo Fisher Scientific) and conjugated with hematoxylin to identify nuclei. Staining was visualized on a Nikon Eclipse microscope at 20X magnification and analyzed using ImageJ from 10 fields per heart.

*Cardiac RNA-Seq*. High throughput sequencing of LV was performed at the University of Missouri DNA Core Facility. Briefly, LV tissue was homogenized in a Tissue-Lyser (Qiagen) and total RNA was extracted using the Qiagen RNeasy Lipid Tissue kit and quantified using a Nanodrop spectrophotometer (Thermo Scientific). Libraries were constructed following the manufacturer's protocol using the Illumina TruSeq mRNA stranded sample preparation kit. The RNA input concentration was determined using the Qubit HS RNA assay kit and Qubit flourometer (Invitrogen) and the RNA quality assessed using the Fragment Analyzer automated electrophoresis system (Agilent). Briefly, the poly-A containing mRNA is purified from total RNA, fragmented, double-stranded cDNA is generated from fragmented RNA, the index containing adapters are ligated, and the amplified cDNA constructs were purified by addition of AxyPrep Mag PCR Clean-up beads. The final construct of each purified library was evaluated using the Fragment Analyzer, quantified using the Qubit HS dsDNA assay kit and fluorometer, and diluted according to Illumina's standard sequencing protocol for sequencing on the NextSeq 500 via single end 75 base pair reads.

RNA-Seq data was processed and analyzed as previously described[9]. Briefly, latent Illumina adapter sequence is identified and removed from input 100-mer RNA-Seq data using Cutadapt. Subsequently, input RNA-Seq reads are trimmed and filtered to remove low quality nucleotide calls and whole reads, respectively, using the Fastx-Toolkit. To generate the final set of quality-controlled RNA-Seq reads, foreign or undesirable sequences are removed by similarity matching to the Phi-X genome (NC_001422.1), the relevant ribosomal RNA genes as downloaded from the National Center for Biotechnology Information, or repeat elements in RepBase, using Bowtie. This final set of quality-controlled RNA-Seq reads is aligned to the Ensemble *Mus musculus* genome sequence, GRCm38.p5, using STAR with the default settings, which also generates the initial expression estimates for each annotated gene. The R Bioconductor package DESeq2 is used to normalize the gene expression estimates across the samples and to analyze the differential expression of genes between sample types. Potential outlier samples are identified and removed at this stage using a combination of Principle Component Analysis and the R libraries nortest and outliers. Gene expression estimates are recalculated after outliers are removed. A gene is identified as being differentially expressed between two conditions when the FDR-corrected p-value of its expression ratio is less than 0.05. Subsequent data is reformatted, sorted and filtered using a variety of R commands and Bash command-line scripts, which are available upon request. Ingenuity Pathway Analysis (IPA; Qiagen) was utilized for examination of the top differentially up- and down-regulated genes and the corresponding top networks, pathways, and associated biological processes.

*Cardiac non-cardiomyocyte single cell preparation.* Single cell suspensions from isolated mouse hearts were prepared, as previously described[10]. Briefly, mice were euthanized with isoflurane and the heart exposed via bilateral thoracotomy before perfusion with DPBS (0.8 mM $CaCl_2$; 10 min). Hearts were subsequently isolated, the atria, valves, and right ventricle removed, and the LV minced to ~1 mm cubes. Minced LV tissue was digested in perfusion buffer containing collagenase IV (2 mg/ml; Worthington Biochemical) and Dispase II (1.2 units/ml; Sigma-Aldrich) for 45 mins at 37°C with suspension trituration every 15 min using 1000 µl micropipettes. The resulting cell suspension was filtered through a 70 µm filter, diluted in 15 ml perfusion buffer, and pelleted at 200 rcf for 20 mins at 4°C with centrifuge brakes disengaged. Cell supernatant was aspirated, the pellet resuspended in perfusion buffer, re-pelleted as described above. The resulting cell pellets were resuspended in FACS buffer (HBSS, 2% FBS; Gibco), passed through a 40 µm filter, pelleted as described above, and resuspended in FACS buffer for subsequent staining and cell sorting. Intact, nucleated non-myocyte cells were subsequently isolated via flow cytometry after staining with Vybrant DyeCycle Ruby nuclear stain (10 µM; ThermoFisher V10273) and SYTOX Green viability stain (30 nM; ThermoFisher S7020).

*Single-cell RNA library preparation and sequencing.* Libraries were constructed by following the manufacturer's protocol with reagents supplied in 10x Genomics Chromium Next GEM Single Cell 3′ Kit v3.1. Briefly, cell suspension concentration and viability were measured manually and with an Invitrogen Countess II automated cell counter. Cell suspension (900 cells per microliter), reverse transcription master mix, and partitioning oil were loaded on a Chromium Next GEM G chip with a cell capture target of 5,000 cells per library. Post-Chromium controller GEMs were transferred to a PCR strip tube and reverse transcription performed on an Applied Biosystems Veriti thermal cycler at 53°C for 45 minutes. cDNA was amplified for 12 cycles and purified using Axygen AxyPrep MagPCR Clean-up beads. cDNA fragmentation, end-repair, A-tailing and ligation of sequencing adaptors was performed according to manufacturer specifications. The final library was quantified with the Qubit HS DNA kit and the fragment size was analyzed using an Agilent Fragment Analyzer system. Libraries were pooled and sequenced on an Illumina NovaSeq to generate 50,000 reads per cell with a sequencing configuration of 28 base pair (bp) on read1 and 98 bp on read2. Isolation of single cells was performed in three batches on separate days with samples from each treatment group included each day to mitigate batch effects.

*Analysis of single-cell RNA-Seq data.* The raw sequencing data were processed using Cell Ranger version 3.1.0 (10x Genomics) before the subsequent analysis. Cell Ranger pipeline used fastq files and aligned sequencing reads to the mm10 transcriptome version 3.0.0 to quantify the expression of genes in each cell. This resulted in data for 83,669 cells that passed quality control steps implemented in Cell Ranger. The filtered count data matrices obtained from cell ranger software were then used for the subsequent analysis. Analyses of scRNA-seq processed data were performed in R version 3.6 and 4.0.1[11] using Seurat suite versions 3.0[12] and tidyverse[13] packages. Further quality control measures of cells with <200 or >8000 expressed genes and genes that were expressed in less than 3 cells were applied per sample manner. Also, cells with more than 30% of UMI mapping to mitochondrial genes were filtered out to control dead or damaged cells. These steps further removed 298 cells from the analysis. The final dataset contains 83,371 cells from 12 mice in 4 conditions and gene expression information for 19,905 genes.

Dimensionality reduction was performed using principal component analysis (PCA) to explore transcriptional heterogeneity and clustering. PC loading for 40 PCs were used as input for a graph-based clustering approach to cluster cells with clustering resolution 0.8. Cells and clusters were visualized on a t-distributed stochastic neighbor embedding (t-SNE) two-dimensional plot generated using the same PC loadings used for the clustering. To optimize the tSNE plot, 1000 iterations and 289 perplexities were used. The identified cell clusters were then annotated based on known marker genes (see Table S8). Figures were primarily generated using Seurat and ggplot2 R packages[14].

*Differential expression analysis.* The differential expression (DE) analysis was performed for each cell population separately. To identify DE genes between groups, we first identified genes expressed in at least 10% of cells in at least one of the groups being compared. We then used MAST R package version 1.12.0[15] to perform DE testing method MASTcpmDetRate considering the cellular detection rate as a covariate. A threshold of uncorrected p <0.01 was used to define statistically significant DE genes between groups.

*Gene ontology analysis.* Gene Ontology (GO) over-representation analysis for differentially expressed gene lists (uncorrected p<0.01) was performed using the enrichGO function from clusterProfiler R package version 3.16.1[16]. The R package org.Mm.eg.db: Genome wide annotation for Mouse, R package version 3.11.4[17] was used to obtain all gene ontology mappings. The over-representation of GO Biological Process terms (GO-BP) was calculated using the entire list of genes identified in the experiment as the background gene list for *Mus musculus* with minimum and maximum gene set sizes 10 and 500, respectively. The similarity between enriched GO-BP terms were calculated using the simplify R function from clusterProfiler R package. GO-BP terms with semantic similarity more than 0.7 were treated as redundant terms and discarded from the analysis. The Benjamini-Hochberg adjusted p-value cut-off of 0.05 was used to determine statistically significant GO-BP terms.

*Flow cytometry.* Immune cells were isolated from the heart by enzymatic digestion, as previously described[18]. Briefly, hearts were isolated, flushed with HBSS, and manually digested into ~1 $mm^3$ pieces. Heart pieces were then enzymatically digested in collagenase II (150 U/mL; Worthington Biochemical) and trypsin (0.6 mg/mL; Worthington Biochemical) at 37°C with agitation. Following digestion, myocyte and non-myocyte fractions were separated by centrifugation at 8x g for 5 min. The non-myocyte containing supernatant was passed through a 70 µm cell strainer prior to flow cytometry analysis.

Cells were stained in 1% FBS in PBS for 30 min at 4°C with the following antibodies: LIVE/DEAD Fixable Aqua Dead Cell Stain Kit (1:40, Invitrogen #L34957), CD3-PE-Cy7 (1:100; Biolegend #100220), CD4-PE (1:100; BD Biosciences #5530449), CD11b-FITC (1:200; Biolegend #101206), CD68-PE (1:50; Biolegend #137014), CD45-BV480 (1:100; BD Biosciences #746682), CD80-PE-Cy7 (1:100; Biolegend #104712), CD117-APC-H7 (1:100; BD Biosciences #560185) and CD196-BV711 (1:100; BD Biosciences #740648). Positive staining was identified based on single antibody controls which were performed for all antibodies on all tissues examined and fluorescence minus one controls were performed on splenic samples to validate cell staining. Isotype controls were also performed on splenic samples using PE-Cy7 rat IgG2b (κ isotype, 1:100, Biolegend 400617), PE rat IgG2a (κ isotype, 1:100, Biolegend 400507), FITC rat IgG2b (κ isotype, 1:100, Biolegend 400633), BV480 rat IgG2b (κ isotype, 1:100, BD Biosciences #565649), BV711 rat IgG2b (κ isotype, 1:100, BD Biosciences #563045) and APC-H7 rat IgG2b (κ isotype, 1:100, BD Biosciences #560200). Following staining, cells were washed twice with PBS and analyzed by flow cytometry using a BD LSRFortessa X-20. Analysis was performed in FlowJo software.

*Cardiac cytokine analysis.* Quantitative proteomic analysis of cardiac cytokines was performed on whole left ventricular lysates by RayBiotech (Mouse Cytokine Array Q4000) and statistical differences between groups were assessed by Wilcoxon test.

*Data Analysis & Statistics*. Data are presented as mean ± standard error with individual data points shown, when appropriate. Statistical analysis was performed using Student *t*-test for planned comparisons, two-way analysis of variance (for repeated measures, when appropriate) with Fisher least significant difference post hoc analysis, as appropriate, in SigmaPlot (SyStat) or Prism (Graphpad). A p value ≤ 0.05 was considered significant.

Supplemental Table 1. *Phenotypic characteristics of study animals by group*

| | Control | Western | Western + Spiro |
|---|---|---|---|
| Body Weight (g) | 22.0 ± 0.7 | 25.7 ± 1.0* | 25.0 ± 0.9* |
| Periovarian Fat Weight (mg) | 431 ± 50 | 834 ± 186* | 790 ± 113* |

Values are mean ± SE, n=7/group; *p<0.05 vs Control.

Supplemental Table 2. *Phenotypic characteristics of study animals by group*

| | MR-Intact Control | MR-Intact Western | SMC MR KO Control | SMC MR KO Western |
|---|---|---|---|---|
| Body Weight (g) | 22.3 ± 0.5 | 29.9 ± 1.0* | 21.5 ± 0.3 | 26.8 ± 1.0*† |
| Heart Weight (mg) | 114 ± 3 | 123 ± 4* | 113 ± 2 | 121 ± 3* |
| HW/TL | 64 ± 2 | 69 ± 2* | 63 ± 1 | 67 ± 1* |
| Periovarian Fat Weight (mg) | 740 ± 82 | 2104 ± 147* | 676 ± 57 | 1540 ± 153*† |
| Systolic Blood Pressure (mmHg) | 120 ± 7 | 119 ± 7 | 117 ± 8 | 124 ± 7 |
| Pulse Wave Velocity (cm/s) | 3.8 ± 0.4 | 3.8 ± 0.3 | 3.5 ± 0.2 | 3.9 ± 0.2 |
| *Plasma Parameters* | | | | |
| Blood Glucose (mg/dl) | 156 ± 11 | 183 ± 10* | 137 ± 7 | 191 ± 8* |
| Plasma Insulin (pg/ml) | 172 ± 25 | 266 ± 50** | 116 ± 24 | 319 ± 35* |
| Plasma Cholesterol (mg/dl) | 75 ± 4 | 115 ± 7* | 70 ± 5 | 110 ± 7* |
| Plasma Triglycerides (mg/dl) | 53 ± 6 | 56 ± 2 | 47 ± 5 | 51 ± 9 |
| Plasma Potassium (mEq/ml) | 4.0 ± 0.2 | 3.8 ± 0.1 | 4.0 ± 0.2 | 4.0 ± 0.2 |
| Plasma Aldosterone (pg/ml) | 763 ± 70 | 1159 ± 160* | 800 ± 61 | 1315 ± 152* |
| *Urine Parameters* | | | | |
| Proteinuria (mg/mg creatinine) | 1.3 ± 0.2 | 2.2 ± 0.2* | 1.1 ± 0.1 | 2.3 ± 0.3* |
| BUN (mg/dl) | 2916 ± 398.2 | 4184 ± 538* | 2149 ± 117 | 4222 ± 422* |

Values are mean ± SE, n=5-23; *p<0.05 vs genotype control, **p=0.08 vs genotype control, †p<0.05 vs MR-intact Western

*Supplemental Table 3.* Cardiac function outcomes by experimental group

| Parameter | **Control** | **Western** | **Western + Spiro** |
|---|---|---|---|
| Heart Rate (bpm) | 542 ± 32 | 512 ± 27 | 525 ± 9 |
| *Morphological parameters* | | | |
| SWTd (mm) | 0.82 ± 0.03 | 0.89 ± 0.09 | 0.86 ± 0.04 |
| PWTd (mm) | 0.71 ± 0.03 | 0.81 ± 0.03§ | 0.73 ± 0.05 |
| LVIDd (mm) | 3.58 ± 0.07 | 3.98 ± 0.13* | 3.55 ± 0.12† |
| LVIDs (mm) | 1.98 ± 0.16 | 2.38 ± 0.16 | 2.01 ± 0.16 |
| RWT | 0.43 ± 0.01 | 0.43 ± 0.04 | 0.45 ± 0.02 |
| LA (mm) | 1.65 ± 0.09 | 2.02 ± 0.05* | 1.97 ± 0.12* |
| Ao (mm) | 1.40 ± 0.04 | 1.49 ± 0.09 | 1.48 ± 0.04 |
| LA/Ao | 1.18 ± 0.06 | 1.39 ± 0.10§ | 1.33 ± 0.07 |
| *Diastolic parameters* | | | |
| E ($m{\cdot}s^{-1}$) | 531.4 ± 28.7 | 602.3 ± 14.8 | 639.2 ± 33.8 |
| E' ($m{\cdot}s^{-1}$) | 23.7 ± 2.5 | 19.6 ± 1.9 | 26.4 ± 2.9 |
| A' ($m{\cdot}s^{-1}$) | 21.5 ± 2.0 | 25.5 ± 1.9 | 27.8 ± 2.0 |
| E/E' | 23.4 ± 2.8 | 32.3 ± 3.3* | 22.2 ± 1.0† |
| E'/A' | 1.17 ± 0.10 | 0.77 ± 0.06* | 0.94 ± 0.06*‡ |
| IVRT (ms) | 12.7 ± 0.9 | 12.7 ± 0.9 | 13.0 ± 0.9 |
| *Systolic parameters* | | | |
| EF (%) | 75 ± 4 | 71 ± 3 | 76 ± 3 |
| FS (%) | 44 ± 3 | 40 ± 2 | 45 ± 3 |

SWTd, septal wall thickness-diastole; PWTd, posterior wall thickness-diastole; RWT, relative wall thickness; LVIDd, LV inner dimension-diastole; LVIDs, LV inner dimension-systole; EF, ejection fraction; FS, fractional shortening; LA, left atrial diameter; Ao, aortic diameter; A', peak late septal annular velocity; E', early peak septal annular velocity; IVRT, isovolumic relaxation time; E, velocity of early mitral inflow; E/E' index of LA filling pressure; MPI, myocardial performance index. Values are mean ± SE, N=4-6. *$p<0.05$ versus Control; †$p<0.05$ versus Western Diet; ‡$p=0.08$ versus Western Diet; §$p=0.07$ versus Control.

*Supplemental Table 4.* Cardiac function outcomes by experimental group

| Parameter | **MR-Intact Control** | **MR-Intact Western** | **SMC MR KO Control** | **SMC MR KO Western** |
|---|---|---|---|---|
| Heart Rate (bpm) | 423 ± 12 | 424 ± 14 | 406 ± 18 | 434 ± 16 |
| *Morphological parameters* | | | | |
| SWTd (mm) | 0.77 ± 0.03 | 0.91 ± 0.03* | 0.70 ± 0.03 | 0.81 ± 0.03‡ |
| PWTd (mm) | 0.68 ± 0.02 | 0.77 ± 0.03* | 0.70 ± 0.04 | 0.74 ± 0.03 |
| LVIDd (mm) | 3.51 ± 0.09 | 3.47 ± 0.08 | 3.49 ± 0.10 | 3.49 ± 0.08 |
| LVIDs (mm) | 2.37 ± 0.12 | 2.23 ± 0.07 | 2.41 ± 0.10 | 2.30 ± 0.10 |
| RWT | 0.42 ± 0.02 | 0.49 ± 0.01* | 0.40 ± 0.02 | 0.45 ± 0.02‡ |
| LA (mm) | 1.59 ± 0.06 | 1.74 ± 0.09 | 1.40 ± 0.07 | 1.53 ± 0.06† |
| Ao (mm) | 1.43 ± 0.09 | 1.32 ± 0.03 | 1.35 ± 0.09 | 1.31 ± 0.03 |
| LA/Ao | 1.14 ± 0.08 | 1.33 ± 0.07* | 1.05 ± 0.05 | 1.18 ± 0.04† |
| *Diastolic parameters* | | | | |
| E ($m\cdot s^{-1}$) | 477.1 ± 18.2 | 551.9 ± 16.9* | 539.2 ± 17.4 | 571.6 ± 22.9 |
| E' ($m\cdot s^{-1}$) | 19.5 ± 0.8 | 18.5 ± 1.4 | 21.0 ± 1.4 | 23.1 ± 1.9† |
| A' ($m\cdot s^{-1}$) | 16.9 ± 0.8 | 19.9 ± 1.1§ | 19.0 ± 0.9 | 20.1 ± 1.3 |
| E/E' | 25.2 ± 0.8 | 30.6 ± 1.9* | 26.3 ± 1.5 | 24.9 ± 1.6† |
| E'/A' | 1.20 ± 0.04 | 0.86 ± 0.05* | 1.12 ± 0.07 | 1.17 ± 0.08† |
| IVRT (ms) | 15.4 ± 0.7 | 16.8 ± 1.0 | 15.1 ± 1.1 | 14.2 ± 0.8 |
| *Systolic parameters* | | | | |
| EF (%) | 61 ± 2 | 66 ± 2§ | 60 ± 2 | 62 ± 2 |
| FS (%) | 33 ± 2 | 35 ± 2 | 31 ± 1 | 33 ± 1 |

SWTd, septal wall thickness-diastole; PWTd, posterior wall thickness-diastole; RWT, relative wall thickness; LVIDd, LV inner dimension-diastole; LVIDs, LV inner dimension-systole; EF, ejection fraction; FS, fractional shortening; LA, left atrial diameter; Ao, aortic diameter; A′, peak late septal annular velocity; E′, early peak septal annular velocity; IVRT, isovolumic relaxation time; E, velocity of early mitral inflow; E/E' index of LA filling pressure; MPI, myocardial performance index. Values are mean ± SE, N=9-14. *$p<0.05$ versus MR-Intact Control; †$p<0.05$ versus MR-Intact Western; ‡$p<0.05$ versus SMC MR KO Control; §$p=0.07$ versus MR-Intact Control.

Supplemental Table 5. *Diameters of isolated coronary vessels by experimental group*

| | Control | Western | Western + Spiro | |
|---|---|---|---|---|
| Diameter (µm) | 243 ± 4 | 257 ± 9 | 256 ± 6 | |
| | **MR-Intact Control** | **MR-Intact Western** | **SMC MR KO Control** | **SMC MR KO Western** |
| Diameter (µm) | 201 ± 10 | 226 ± 8 | 214 ± 7 | 212 ± 8 |

Values are mean ± SE, n=5-14; Diameter calculated from circumference following the normalization procedure.

*Supplemental Table 6.* Primer sequences for real-time quantitative PCR

| Gene Name | Primer Sequence (5' → 3') Forward | Reverse |
|---|---|---|
| **ITGAX (CD11c)** | CTGGATAGCCTTTCTTCTGCTG | GCACACTGTGTCCGAACTC |
| **ADGRE1 (F4/80)** | CTTTGGCTATGGGCTTCCAGTC | GCAAGGAGGACAGAGTTTATCGTG |
| **GAPDH** | TCACCACCATGGAGAAGGC | GCGAAGCAGTTGGTGGTGCA |
| **ICAM-1** | AACCGCCAGAGAAAGATCAG | TGTGACAGCCAGAGGAAGTG |
| **PECAM-1** | GAGCCCAATCACGTTTCAGTTT | TCCTTCCTGCTTCTTGCTAGCT |
| **TNF-α** | GCCTCTTCTCATTCCTGCTTG | CTGATGAGAGGGAGGCCATT |
| **VCAM-1** | CTTCATCCCCACCATTGAAG | TGAGCAGGTCAGGTTCACAG |

**Supplemental Tables 7-16 are available at the following repository:**

https://github.com/pinto-lab/Dona-et-al_2022_SMC-MR-KO_obesity

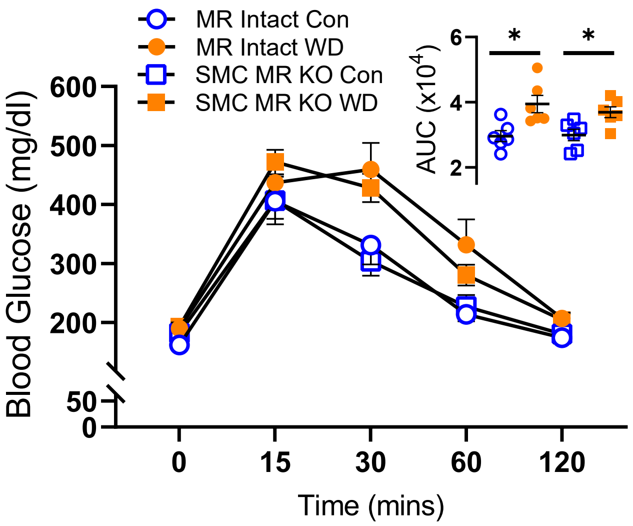


**Supplemental Figure 1. Smooth muscle cell mineralocorticoid receptor knockout (SMC MR KO) does not prevent western diet (WD)-induced glucose intolerance in female mice.** Glucose excursions over time during a glucose tolerance test (ip, 1 g/kg) and glucose area under the curve (AUC; inset) in MR Intact and SMC MR KO mice fed control (Con) and WD for 16 weeks. Values are mean±SE with individual data points shown (inset), n=6/group, *p<0.05 for indicated comparison.

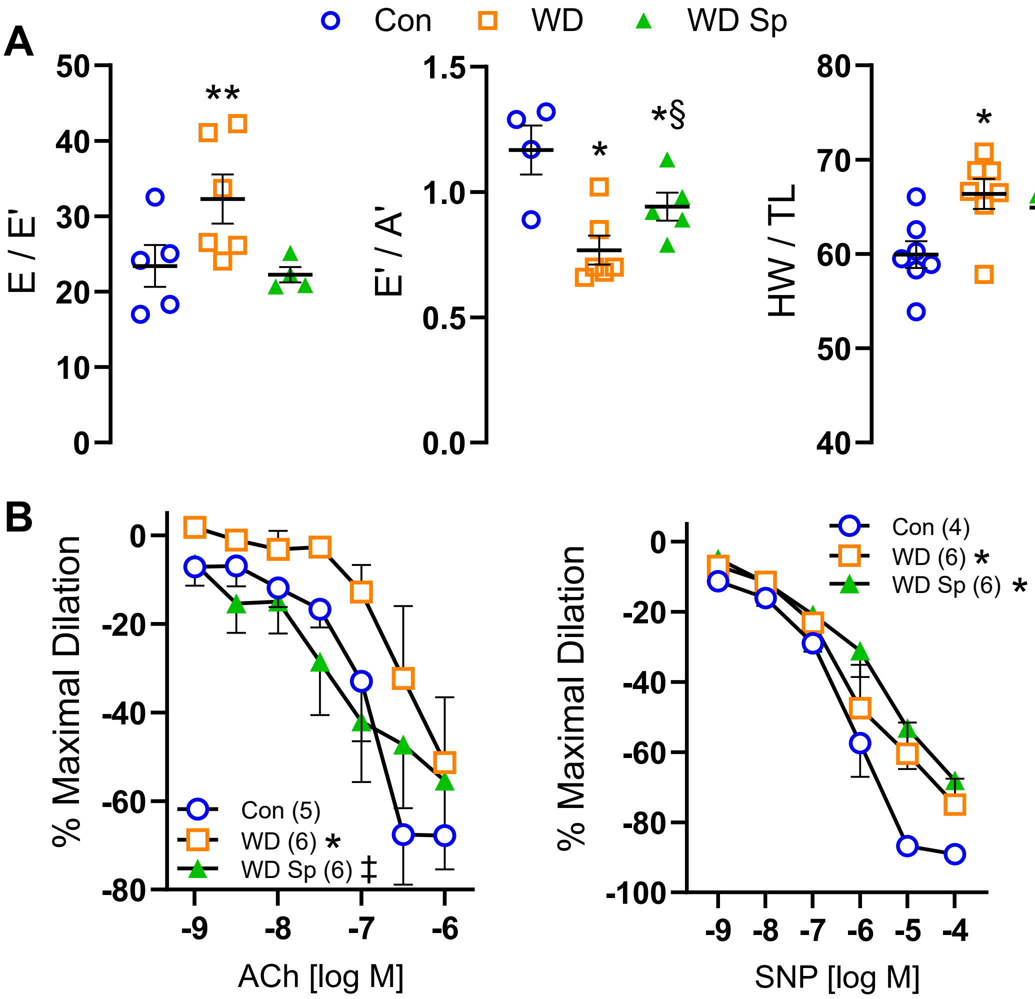


**Supplemental Figure 2. Systemic mineralocorticoid blockade with spironolactone (Sp) prevents western diet (WD)-induced cardiac and coronary vascular dysfunction in female mice.** (A) Indices of cardiac diastolic function, specifically estimated left ventricular filling pressure (E/E') and early-to-late diastolic septal annulus motion ratio (E'/A'), and cardiac weights (heart weight-to-tibia length ratio; HW/TL) in Con and WD fed mice. (B) Vasodilator responses of isolated coronary arteries to endothelium-dependent (acetylcholine, ACh) and -independent (sodium nitroprusside, SNP) agonists. Values are mean±SE with individual data points shown (A) and sample size indicated in parentheses (B); *p<0.05 versus Con, **p<0.05 versus all other groups, ‡p<0.05 versus WD ,§p=0.08 versus WD.

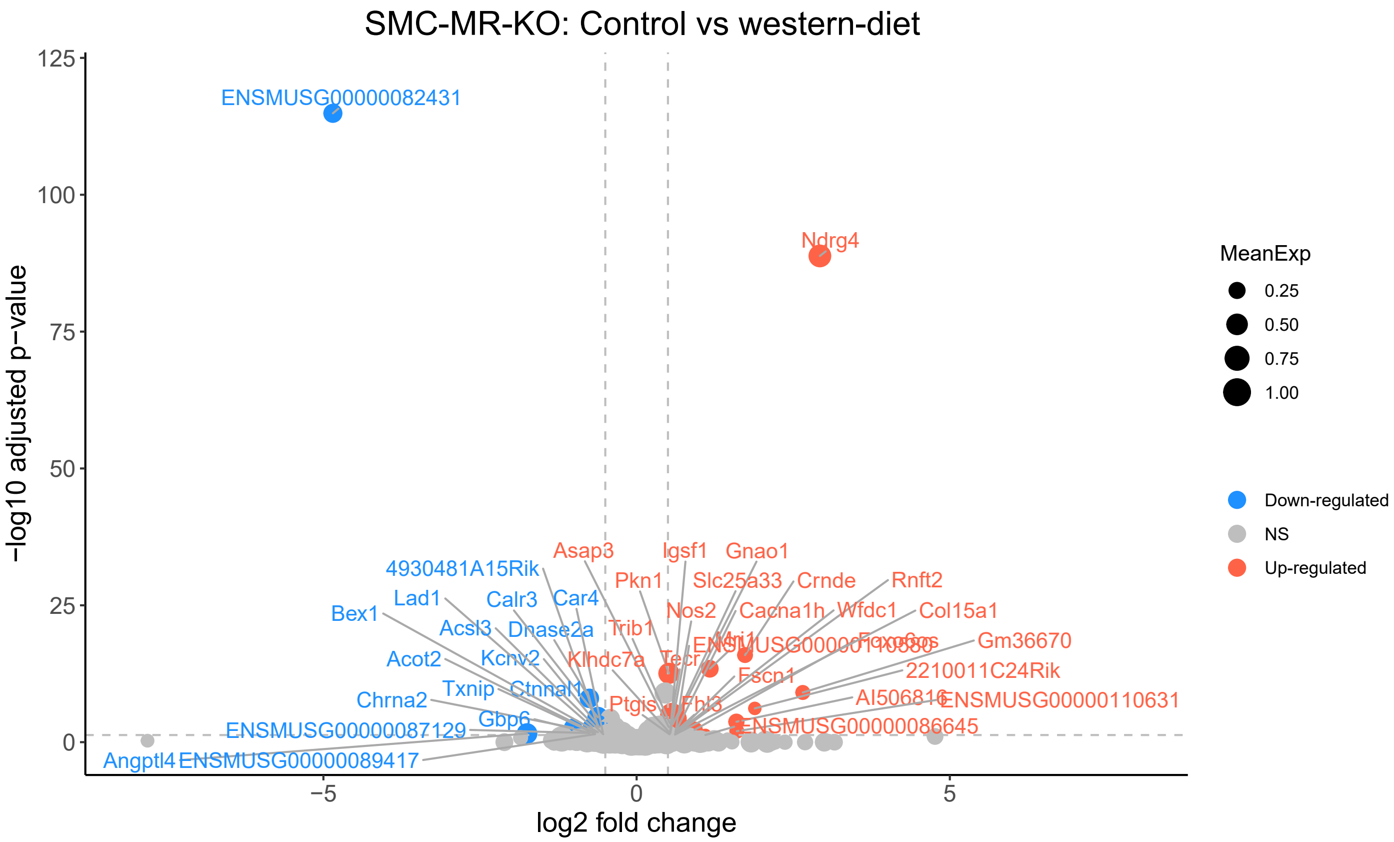


**Supplemental Figure 3. Western diet (WD) feeding altered cardiac gene expression in smooth muscle cell mineralocorticoid receptor knockout (SMC-MR-KO) mice.** Analysis of cardiac transcripts (13,565 transcripts) revealed differential expression (log2 fold change >0.5, corrected p<0.05) of 43 transcripts induced by WD feeding in SMC-MR-KO mice (17 downregulated, blue dots; 26 upregulated, red dots).

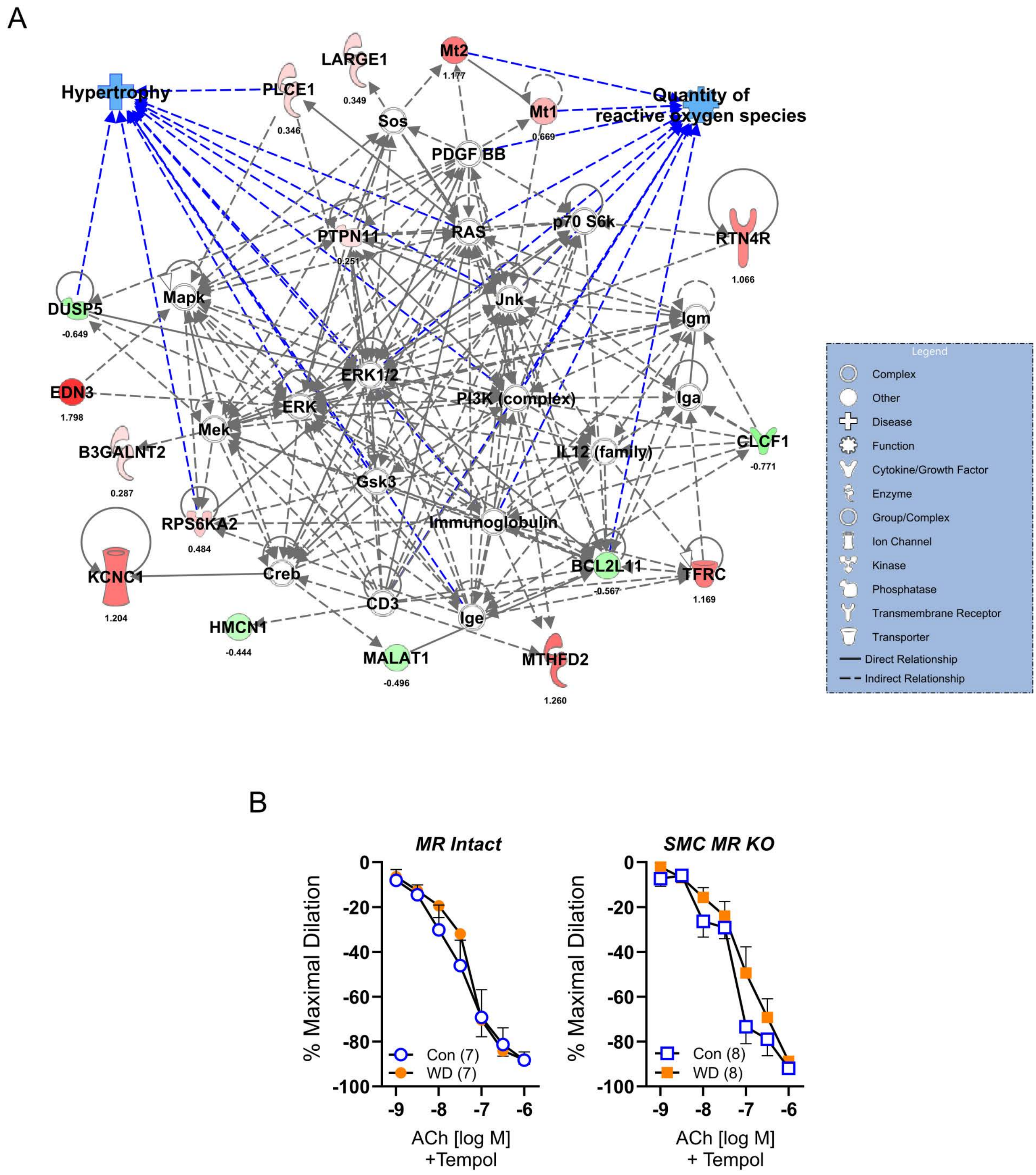


**Supplemental Figure 4. Western diet (WD) feeding alters the cardiac transcriptome in MR Intact mice and induces coronary oxidative stress in female mice that is prevented by smooth muscle cell mineralocorticoid receptor knockout (SMC-MR-KO).** (A) Top differentially regulated IPA network (IPA score=39; green nodes, downregulated; red nodes, upregulated) in WD-fed MR Intact mice versus control (Con)-fed MR Intact mice with log2 fold changes and enriched biological processes indicated. (B) Vasodilator responses of isolated coronary arteries to the endothelium-dependent vasodilator acetylcholine (ACh) in the presence of the superoxide dismutase mimetic Tempol (1 mM). Compare to Figure 1C to view impact of Tempol treatment across groups, especially in vessels from WD-fed MR Intact mice. Values are mean±SE, sample size in parentheses.

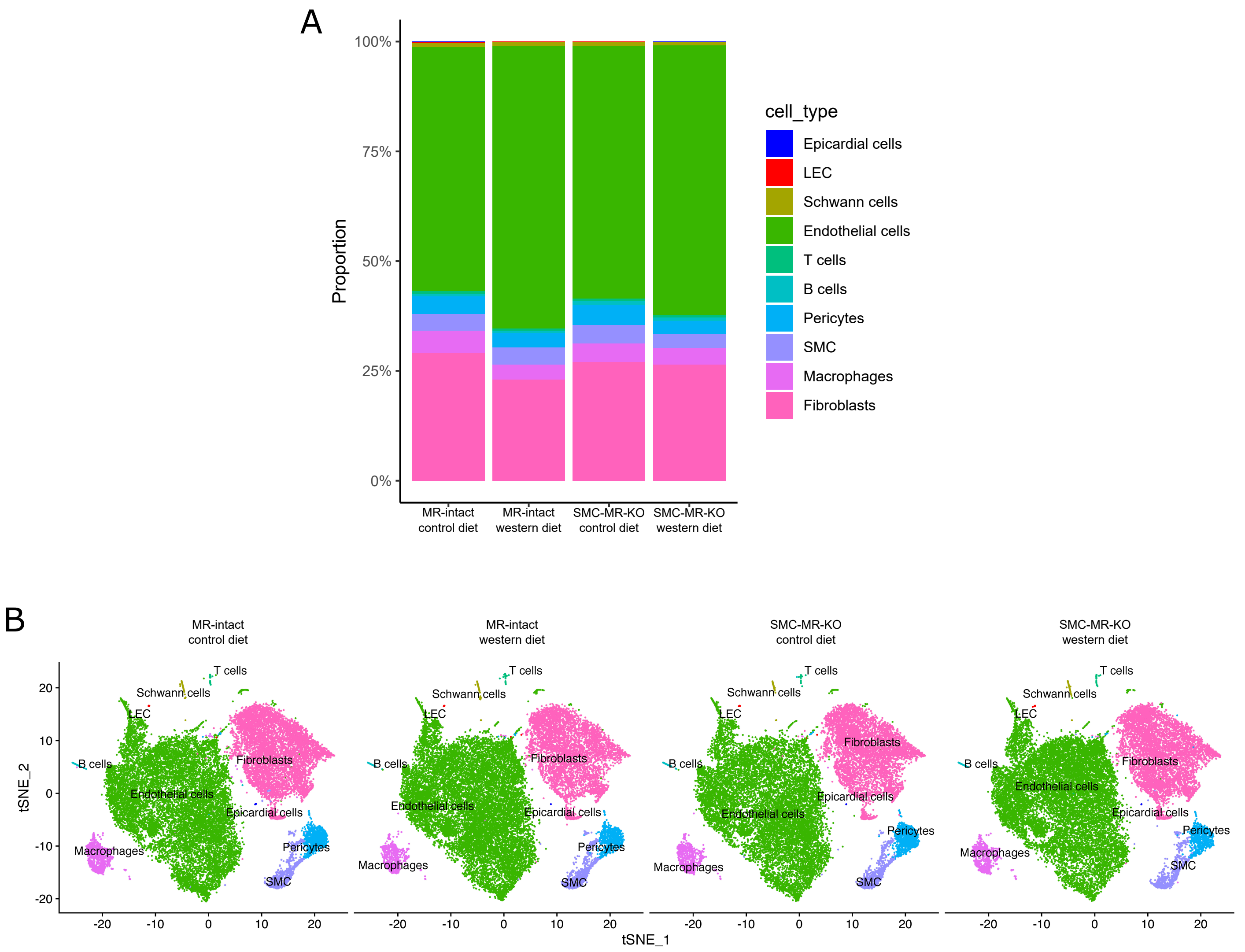


**Supplemental Figure 5. Isolation of cells for single-cell RNA sequencing capture similar proportions of diverse cardiac non-myocyte cell populations across experimental groups.** (A) Relative proportions of cell types analyzed from individual samples from mineralocorticoid receptor (MR) Intact and smooth muscle cell MR knockout (SMC-MR-KO) mouse hearts (n=3 per group). Heights of the individual boxes comprising each bar represent the proportion of cell classified as each cell population. (B) t-SNE projections of all cardiac non-myocyte cells from each experimental group.

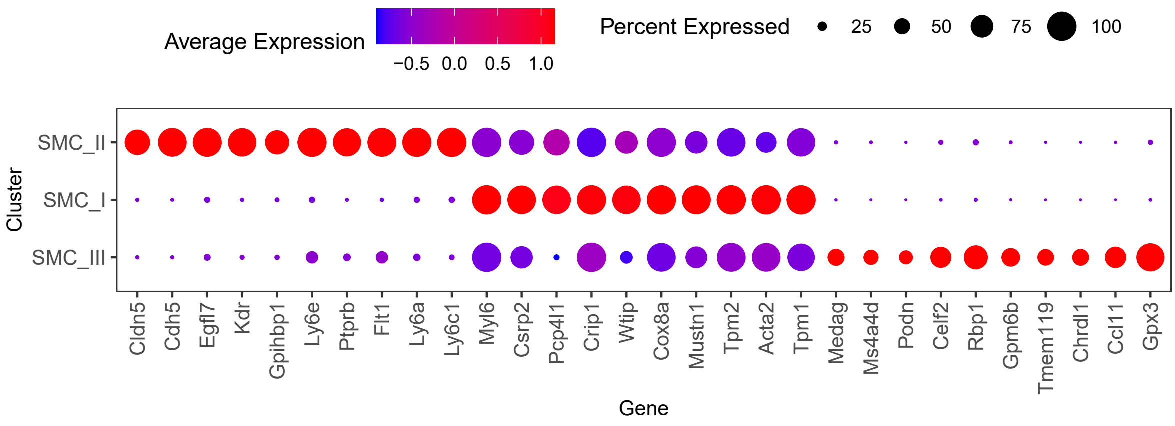


**Supplemental Figure 6. Key gene expression differences between smooth muscle cell (SMC) populations.** Dot plot for top highly and uniquely expressed genes in each smooth muscle cell population identified using an unsupervised analysis. Dot color and size indicate the relative expression and percentage of cells expressing that gene within each cell population, respectively.

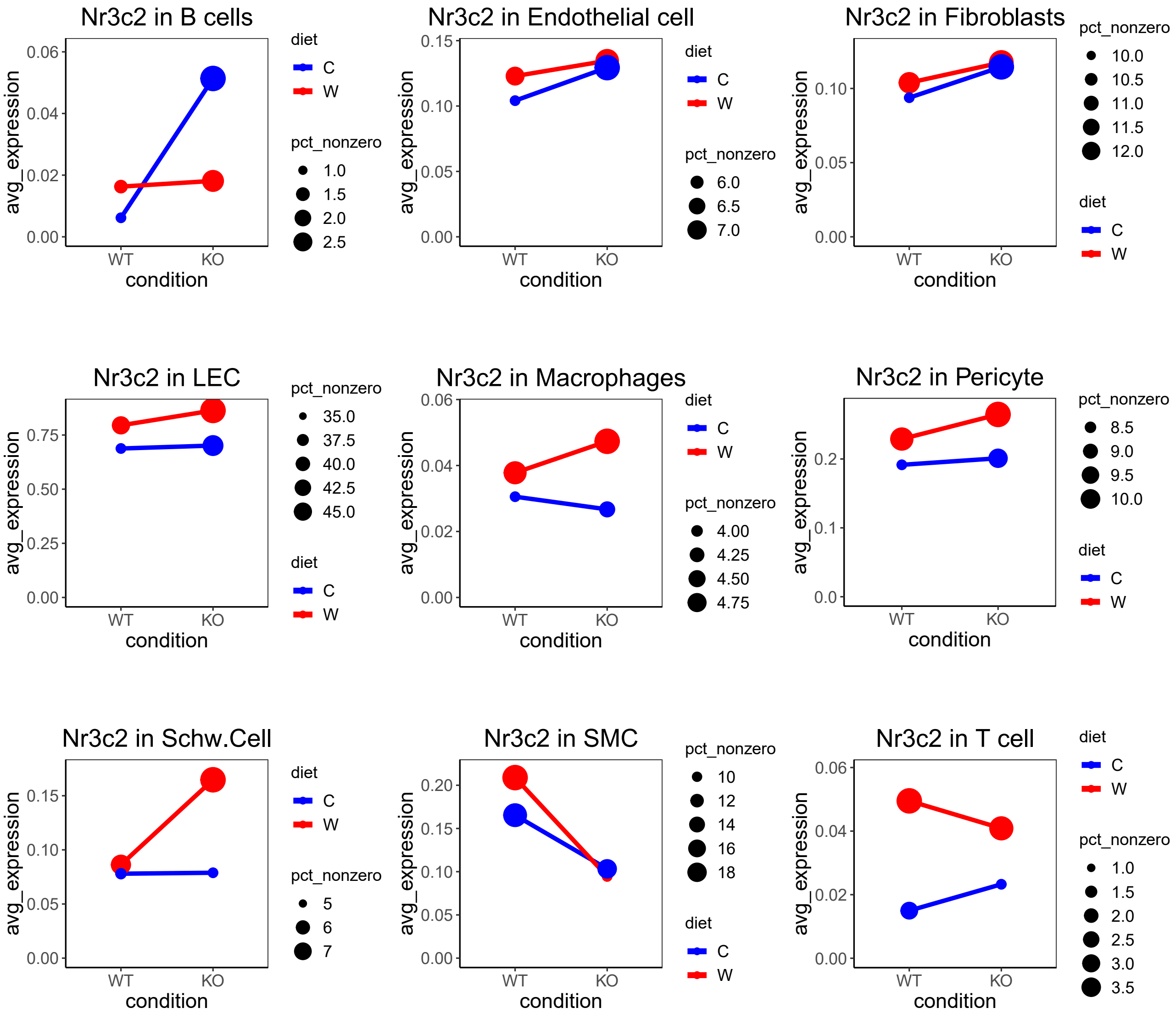


**Supplemental Figure 7. Mineralocorticoid receptor (MR) gene expression in cardiac non-myocyte cell populations with and without western diet.** Gene expression of MR (encoded by Nr3c2) in cardiac non-myocytes from control (C) and WD fed (W) MR Intact (WT) and smooth muscle cell MR knockout (KO) mice. Dot color and size indicate the diet group and the percentage of cell expressing Nr3c2 gene within each group, respectively.

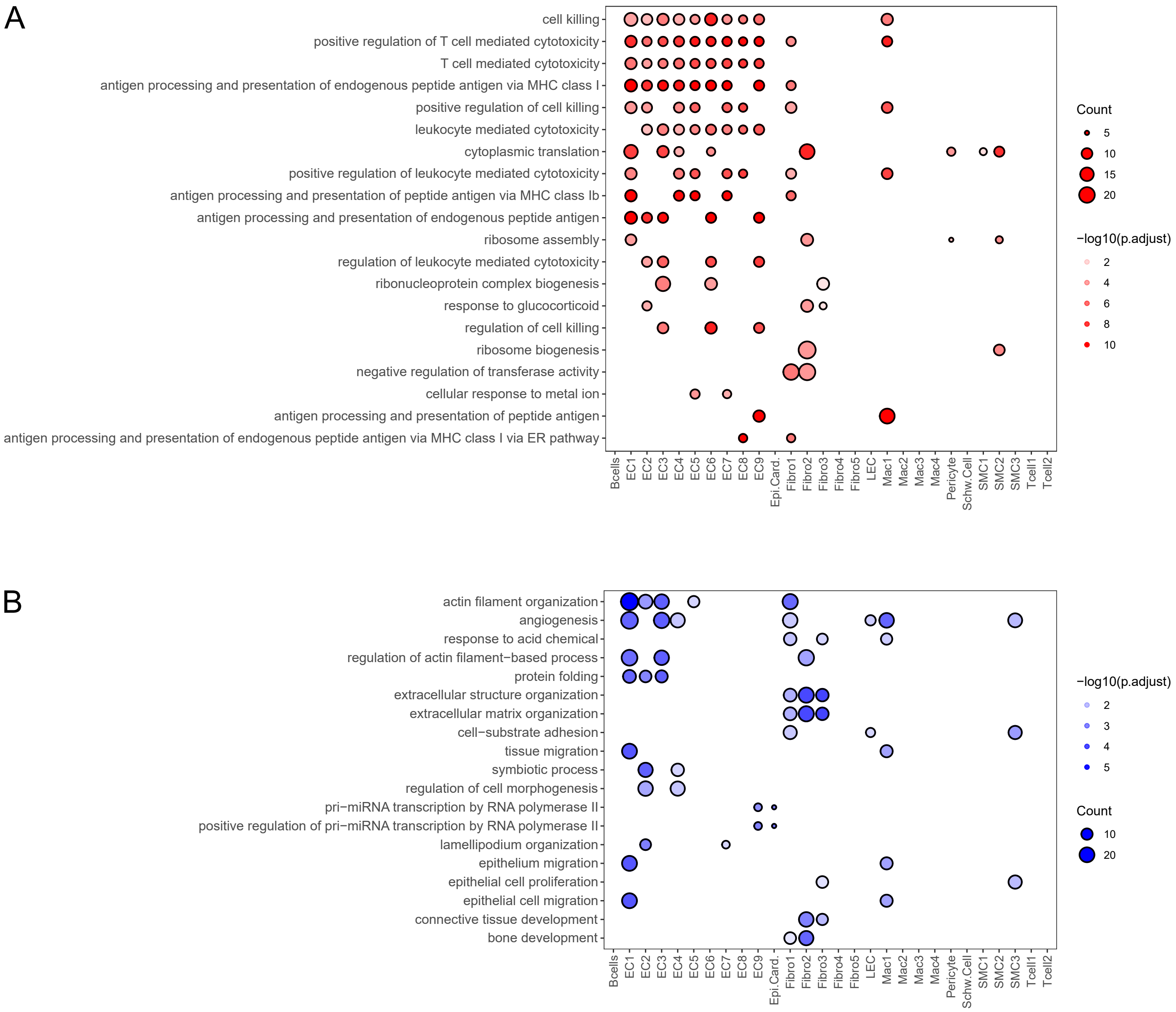


**Supplemental Figure 8. Smooth muscle cell mineralocorticoid receptor knockout (SMC MR KO) alters gene programs across cardiac non-myocyte cell populations.** Dot plots summarizing statistically significant gene ontology (GO) terms (corrected p<0.05) enriched among genes differentially up- (A) or down- (B) regulated in each cell population of control diet fed SMC MR KO mice versus control diet fed MR Intact mice. Dot color intensity and size indicate significance and number of differentially expressed genes related to each GO term, respectively.

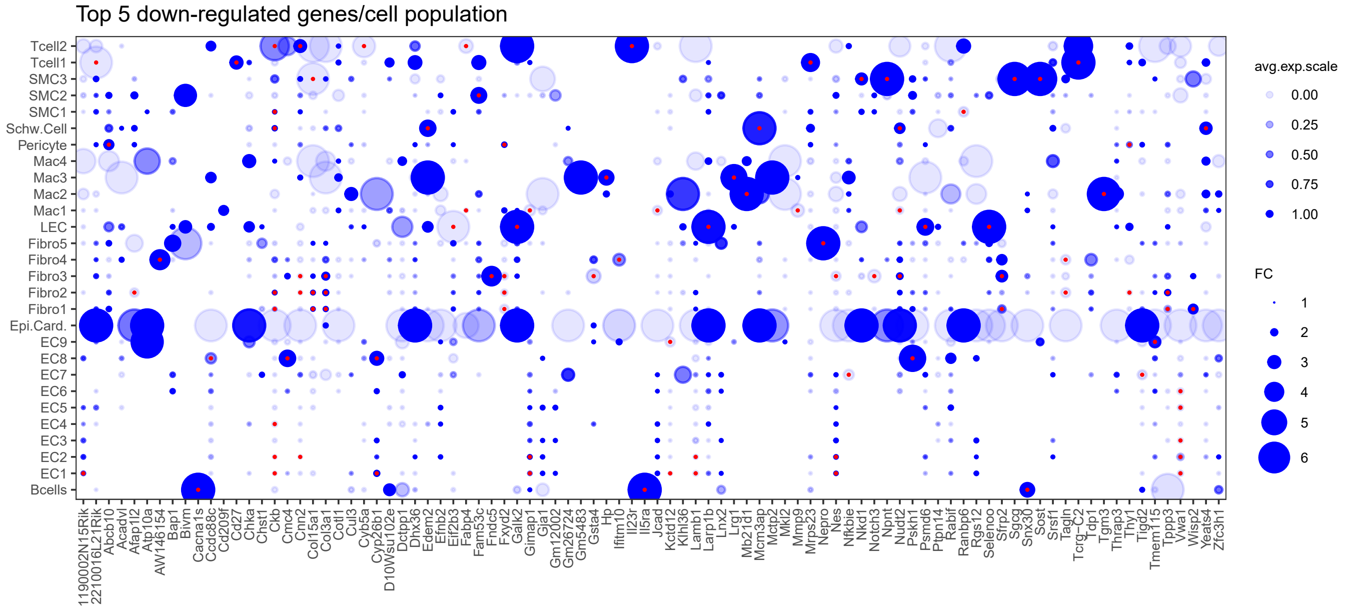


**Supplemental Figure 9. Western diet (WD) feeding down-regulates gene expression in a cell-specific manner in mineralocorticoid receptor (MR) Intact mice.** Dot plot summarizing the relative expression of top significantly down-regulated genes in WD-fed versus control-fed MR Intact mice across cardiac non-myocyte cell populations. Dot color intensity and size are proportional to the relative gene expression in WD cells and the fold change increment in WD cells compared to control cells within each population, respectively. Red points at the centers of some dots highlight statistically significant differences in gene expression in WD cells versus control cells (uncorrected p<0.001).

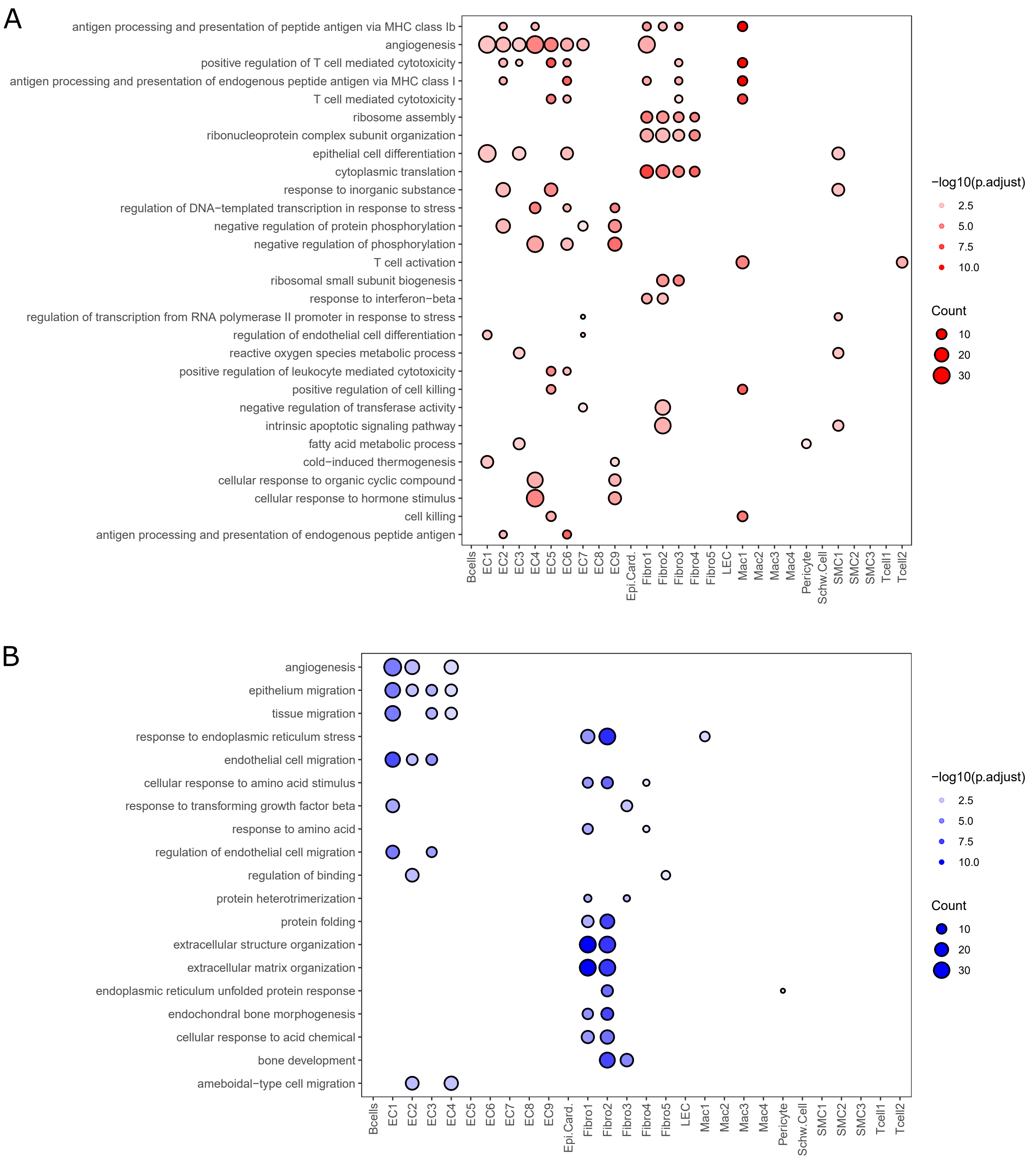


**Supplemental Figure 10. Western diet (WD) feeding impacts gene programs in a cell-specific manner among cardiac non-myocyte populations.** Dot plot summarizing statistically significant gene ontology (GO) terms (corrected $p<0.05$) enriched by WD up- (red dots; A) and down-regulated (blue dots; B) genes in each cell population from MR Intact mice. Dot color intensity and size indicate significance and number of differentially expressed genes related to each GO term, respectively. GO terms displayed and their order within plots is based on frequency of GO terms in difference cell populations.

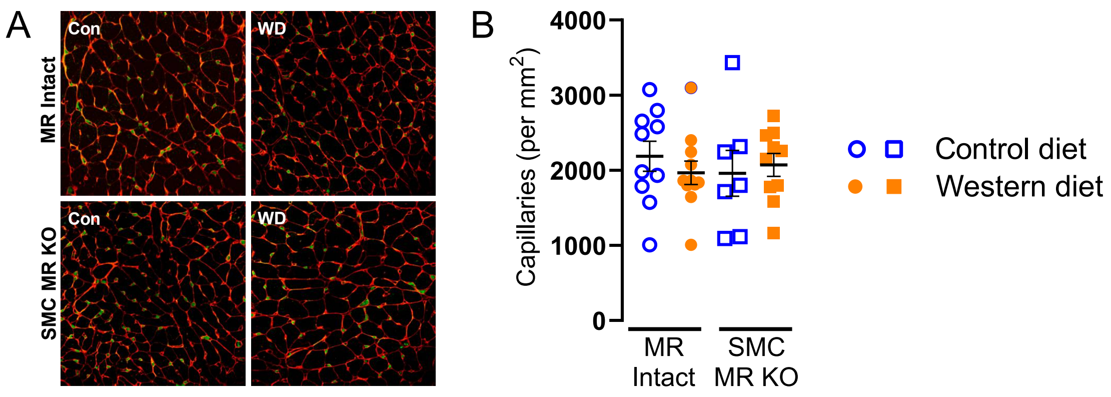


**Supplemental Figure 11. Western diet (WD) or smooth muscle cell mineralocorticoid receptor knockout (SMC MR KO) do not change cardiac capillary density.** (A) Micrographs of cardiac capillary density, assessed by staining endothelial cells (CD31 staining; red) and nuclei (green), in control- and WD-fed MR Intact and SMC MR KO mice. (B) Statistical summary of capillary density derived from micrographs. Values are mean±SE with individual data points shown.

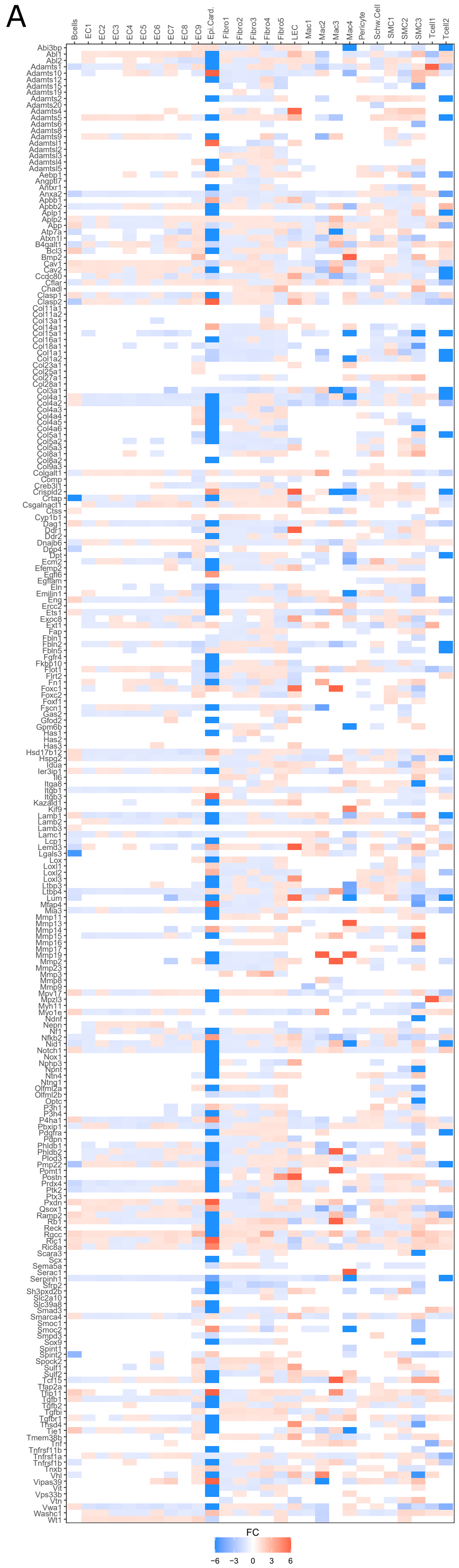


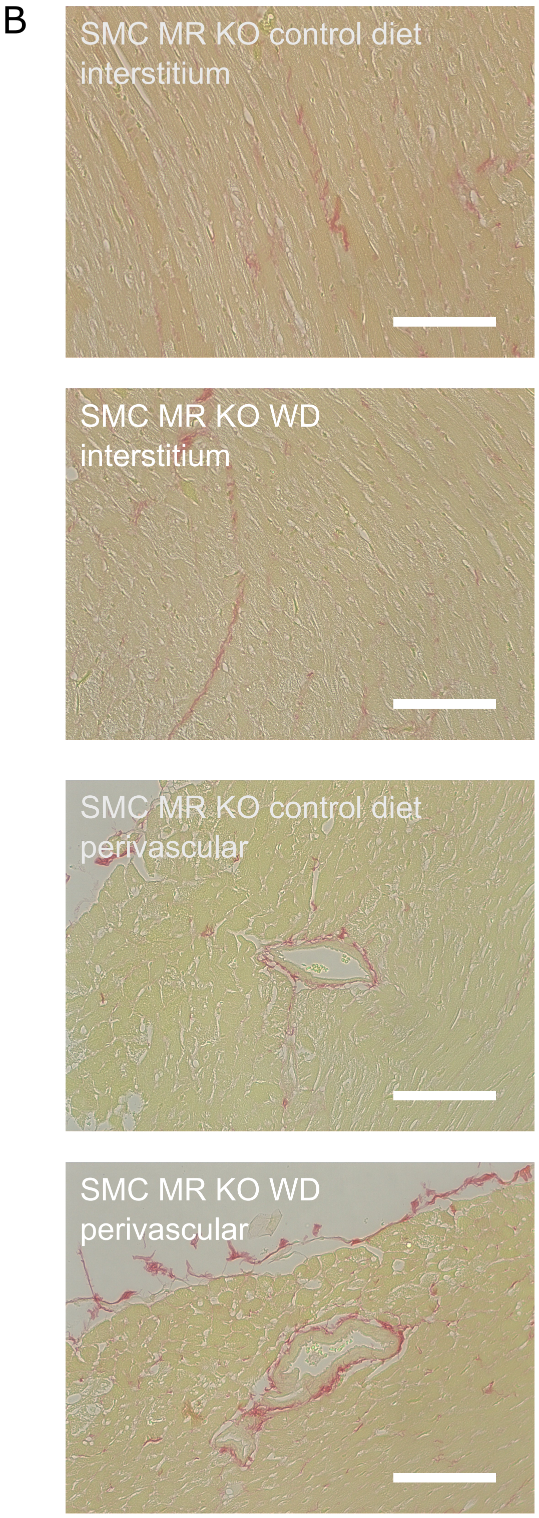


**Supplemental Figure 12. Impact of western diet (WD) feeding on extracellular matrix genes across cardiac non-myocyte cell populations.** (A) Heatmap of extracellular matrix genes (from gene ontology term ‘extracellular matrix organization’, #0030198) and their relative expression in MR intact mice fed either control or WD. Box color and intensity indicate direction: blue indicates down-regulation and red indicates up-regulation of gene after WD. (B) Representative interstitial and perivascular fibrosis, assessed by picrosirius red staining, in control- and WD-fed smooth muscle cell mineralocorticoid receptor knockout (SMC MR KO) mice.

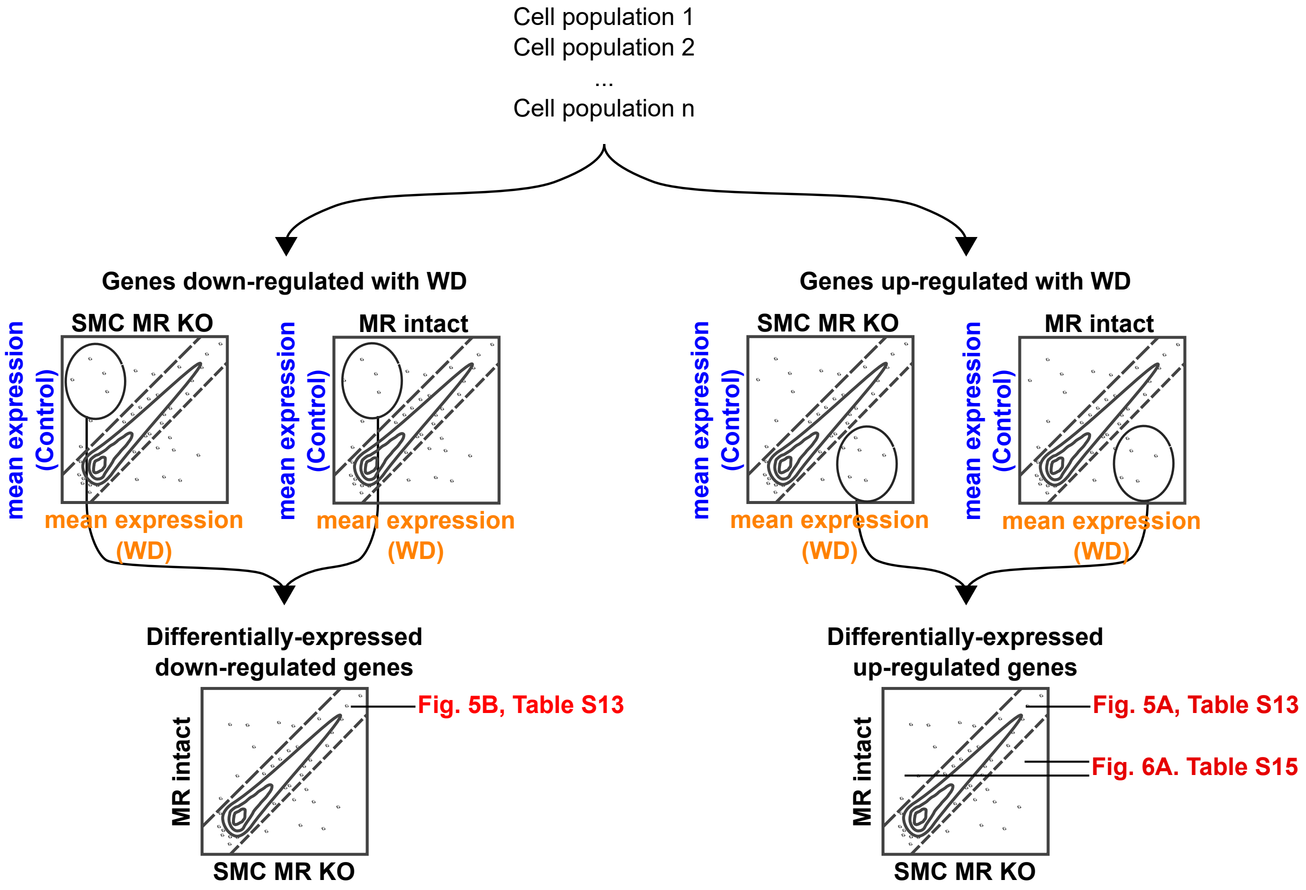


**Supplemental Figure 13. Strategy for determining genes differentially and more robustly impacted by western diet (WD) feeding in mineralocorticoid receptor (MR) Intact versus smooth muscle cell MR knockout (SMC-MR-KO) mice.** For each cell population, MR-Intact and SMC-MR-KO cell populations were considered in isolation for genes upregulated or downregulated after WD. For genotype-specific responses to WD feeding upregulated genes, a list of genes upregulated in either MR-Intact or SMC-MR-KO mice was generated and differences in gene expression examined. For genotype specific responses to WD feeding in downregulated genes, a list of genes downregulated in either genotype was generated and differences in gene expression was again examined.

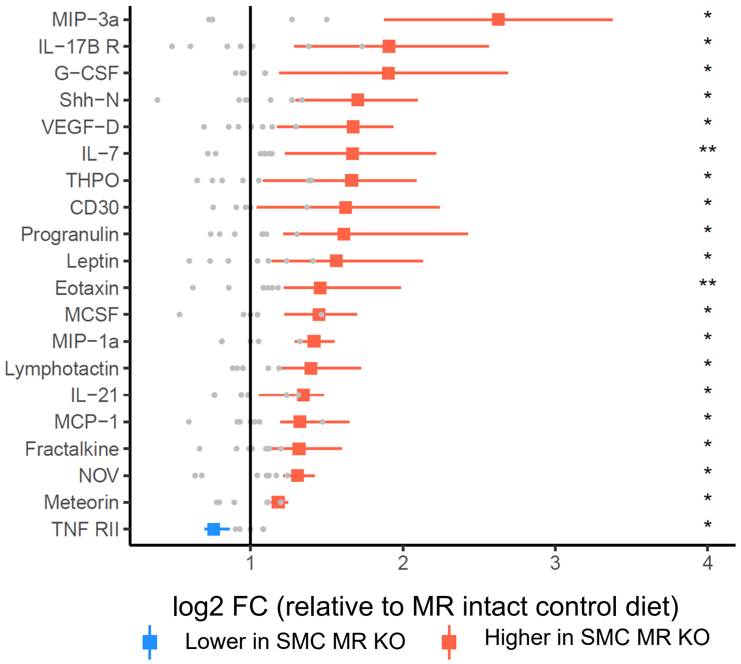


**Supplemental Figure 14. Smooth muscle cell mineralocorticoid receptor knockout (SMC-MR-KO) modulates cardiac cytokines in control diet fed mice.** Differentially expressed cardiac cytokines in control diet fed SMC-MR-KO mice versus control diet fed MR-Intact mice. Black line indicates mean cytokine level in MR-intact mouse hearts. Gray dots indicate individual data points for MR-intact group. Red and blue points indicate cytokines at higher or lower levels (respectively) in SMC-MR-KO hearts, relative to mean of MR-intact mouse hearts. Colored squares a indicate mean values with colored lines indicating value range of data points for SMC-MR-KO samples. *p<0.05, **p<0.01.

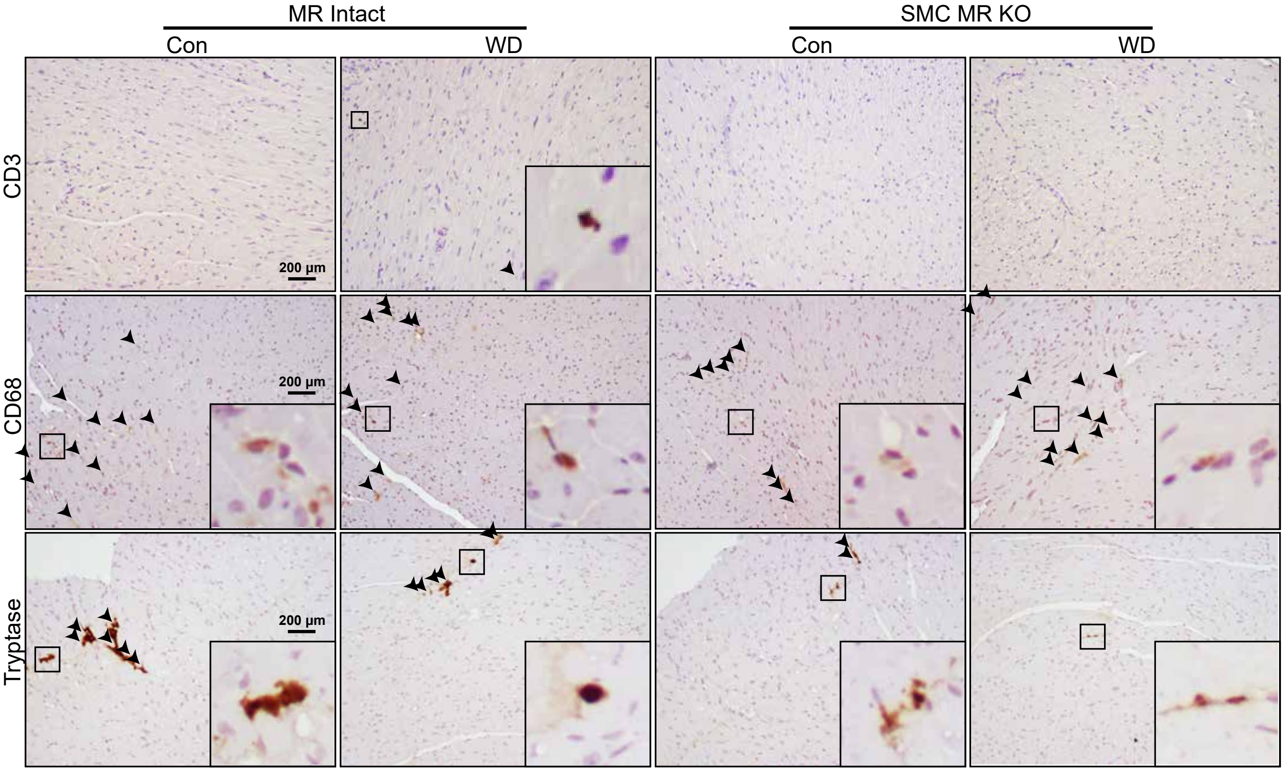


**Supplemental Figure 15. Representative cardiac staining for CD3, CD68, and Tryptase in cardiac sections from mineralocorticoid receptor (MR) Intact and smooth muscle cell MR knockout (SMC MR KO) mice fed control (Con) and western diet (WD).** Arrows point to positively stained cells, insets include zoomed in view of positive staining indicated by the smaller box.

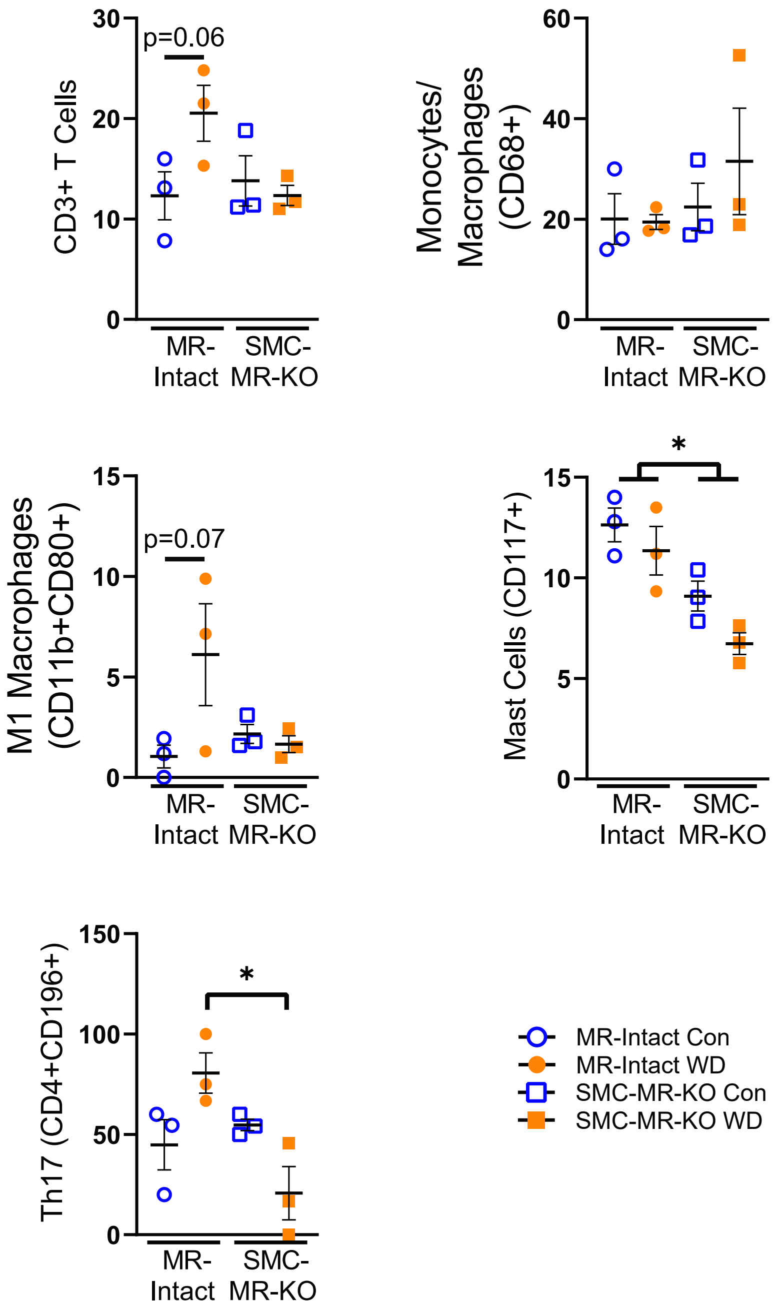


**Supplemental Figure 16. Smooth muscle cell mineralocorticoid receptor knockout (SMC-MR-KO) prevents western diet (WD)-induced cardiac leukocyte recruitment in female mice.** Cardiac leukocytes, specifically CD3+ T cells, CD68+ monocytes/macrophages, CD11b+CD80+ M1 macrophages, CD117+ mast cells, and CD4+CD196+ Th17 cells, assessed by flow cytometry. Values are mean±SE with individual data points shown; *p<0.05 for indicated comparison.
